# Optimizing a CRISPR-Cas13d gene circuit for tunable target RNA downregulation with minimal collateral RNA cutting

**DOI:** 10.1101/2024.05.11.593702

**Authors:** Yiming Wan, Christopher Helenek, Damiano Coraci, Gábor Balázsi

## Abstract

The invention of RNA-guided DNA cutting systems has revolutionized biotechnology. More recently, RNA-guided RNA cutting by Cas13d entered the scene as a highly promising alternative to RNA interference to engineer cellular transcriptomes for biotechnological and therapeutic purposes. Unfortunately, “collateral damage” by indiscriminate off-target cutting tampered enthusiasm for these systems. Yet, how collateral activity, or even RNA target reduction depends on Cas13d and guide RNA abundance has remained unclear due to the lack of expression-tuning studies to address this question. Here we use precise expression-tuning gene circuits to show that both nonspecific and specific, on-target RNA reduction depend on Cas13d and guide RNA levels, and that nonspecific RNA cutting from *trans* cleavage might contribute to on-target RNA reduction. Using RNA-level control techniques, we develop new *Multi-Level Optimized Negative-Autoregulated Cas13d and crRNA Hybrid* (MONARCH) gene circuits that achieve a high dynamic range with low basal on-target RNA reduction while minimizing collateral activity in human kidney cells and green monkey cells most frequently used in human virology. MONARCH should bring RNA-guided RNA cutting systems to the forefront, as easily applicable, programmable tools for transcriptome engineering in biotechnological and medical applications.

## INTRODUCTION

The discovery of CRISPR (“clustered regularly interspaced short palindromic repeats”)^1–4^ has revolutionized biotechnology. The use of various CRISPR-associated (Cas) proteins has facilitated relatively straightforward deletion, insertion, and editing of genes, enabling extensive investigations across bacterial, fungal, plant, and mammalian cells^4–6^. Yet, gene expression levels are as critical as protein-coding sequences in determining cellular phenotypes^7–12^. To manipulate gene expression without altering DNA sequences, enzymatically dead Cas (dCas) proteins, fused with silencing and activator domains, were developed^13^. However, these dCas systems, which control gene expression through DNA binding, may suffer from nonspecific binding, propagation of chromatin modifications, and diversion of cellular resources^14–17^. Recently, RNA-cutting CRISPR systems have emerged as promising alternatives that address these issues by cutting the targeted mRNA instead of gene repression by DNA binding^18^. Nonetheless, challenges such as transcriptome-wide “collateral damage” due to nonspecific RNA cutting from *trans* cleavage initiated automatically upon *cis* on-target cleavage have moderated the enthusiasm for these novel systems^19–20^. Despite these drawbacks, Cas13d represents a potentially crucial, modular, targetable, and programmable alternative to existing RNA interference technologies, promising significant advancements in both research and clinical settings.

Despite considerable information on Cas13d, it remains poorly understood how the expression levels of Cas13d and the corresponding crRNA affect the specific (on-target) and nonspecific activities of the Cas13d-crRNA complex, and consequently, its target RNA reduction. This gap in knowledge is due to a lack of studies employing genetic systems that fine-tune the expression of Cas13d and crRNA as opposed to binary on/off approaches or crRNA mismatches^21–22^, which do not allow for reversible, tunable, precise expression control in the same cells.

Over the past two decades, we have developed and refined negative feedback (NF) synthetic gene circuits for precise eukaryotic gene expression tuning^12, 23–26^, which have been applied in yeast^23^, Chinese hamster^27–28^, and human cells^12, 24–25^, revealing surprising phenotypic effects associated with fine-tuned gene expression changes. For instance, NF expression tuning has revealed an unexpected non-monotone dependence or “landscape” of cellular invasion versus metastasis activator BACH1 expression^12^. Mammalian NF (mNF) systems have not been applied yet to test whether fine-tuned expression of CRISPR/Cas system components could enhance the specificity, efficacy and tunability of CRISPR/Cas action, in particular for CRISPR-Cas13d. Therefore, there is a need and opportunity to map the currently unknown regulatory and fitness landscapes of CRISPR-Cas systems, which requires fine-tuning of their components with synthetic biological sophistication **(Figure 1A)**.

**Figure 1.**
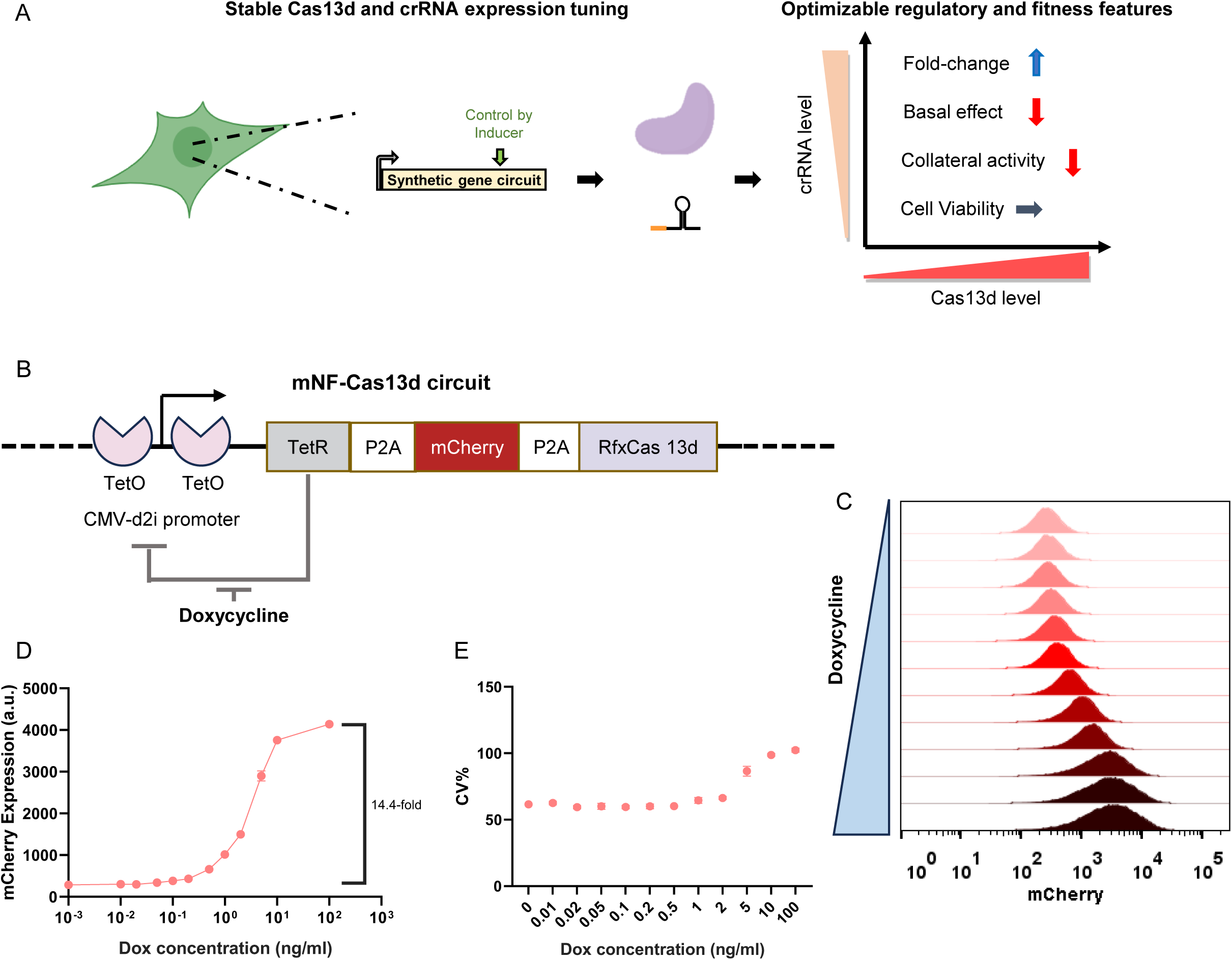
Characterization of the mNF-Cas13d gene circuit LP-integrated into the AAVS1 safe harbor site. **(A)** Illustration of the need to understand and optimize multiple aspects of RNA reduction by various CRISPR/Cas13d RNAse activities. This could be fulfilled by fine-tuning both Cas13d and crRNA expression levels using stably integrated synthetic gene circuits. Arrows indicate a desired optimization of each aspect, where the blue arrow corresponds to an increase, the red arrow corresponds to a decrease and the grey arrow corresponds to no change. **(B)** Diagram of the synthetic mNF-Cas13d gene circuit for Dox-controlled, joint tuning of tetracycline repressor (TetR), mCherry reporter and Cas13d protein levels after AAVS1 site-specific LP-integration. TetO: Tetracycline Operator; P2A: self-cleaving peptide. **(C)** Representative dose-responses of red fluorescence intensity histograms for the stably integrated mNF-Cas13d gene circuit, measured at 0, 0.01, 0.02, 0.05, 0.1, 0.2, 0.5, 1, 2, 5, 10, 100 ng/ml Dox levels, respectively. **(D)** Dose-responses of mean fluorescence intensity of the mCherry reporter for the stably LP-integrated mNF-Cas13d gene circuit in HEK 293 cells (n=3). **(E)** Dose-responses of the coefficient of variation (CV) of mCherry reporter for the stably LP-integrated mNF-Cas13d gene circuit in HEK 293 cells (n=3).

Here, we set out to develop a standard mNF-Cas13d gene circuit that transcriptionally tunes Cas13d expression, to facilitate dose-dependent target RNA reduction in Human Embryonic Kidney (HEK293) cells. Despite the decent dynamic range of Cas13d expression, basal (uninduced) Cas13d expression already reduced target RNA levels substantially, restricting the dynamic range of target RNA downregulation. Furthermore, excessive Cas13d and crRNA expression also depleted non-target reporter RNAs, indicating considerable collateral cutting. To address these limitations, we developed a novel mNF-derived gene circuit design, termed *Multi-Level Optimized Negative-Autoregulated Cas13d and crRNA Hybrid* (MONARCH). This design combines mNF with the inherent ability of Cas13d to process its own crRNA as a mechanism for RNA-level autoregulation^29^, effectively attenuating the expression of both Cas13d and crRNA. Besides eliminating collateral activity, MONARCH achieved a broad dynamic range of both transient and stable RNA target downregulation with minimal basal effect. We further optimized MONARCH by incorporating an mRNA stabilizing Triplex motif, to enable even broader on-target dynamic range without reintroducing collateral activity. Finally, we demonstrated MONARCH’s functionality in Vero E6 green monkey kidney cells, the workhorse of human virology, offering a scalable and versatile cellular platform for quantitative RNA-targeting in antiviral studies and therapeutic development.

## RESULTS

### A versatile genomic landing-pad platform enables rapid and reliable mNF-Cas13d gene circuit integration

To downregulate arbitrary mRNA targets, we set out to precisely tune Cas13d expression using the mNF gene circuit^24^. Thus, we inserted the gene encoding Cas13d as the last component in a 2A-separated multi-gene construct that also contains the repressor hTetR and the fluorescent reporter mCherry. We used D2i, a CMV-derived promoter with *TetO2* binding sites to drive the expression of this hTetR-mCherry-Cas13 construct (mNF-Cas13d gene circuit, **Figure 1B**), ensuring negative autoregulation by hTetR. The expression of all gene circuit components is precisely tunable by adding doxycycline to the growth medium. Doxycycline diffuses into cells, binds hTetR and prevents its DNA binding, elevating mNF-Cas13d expression. Negative autoregulation reduces expression noise for all circuit genes up until saturation, ensuring precise Cas13d gene expression tuning.

To enable subsequent single-copy, harmless, mutation-free, and precise insertion of the mNF-Cas13d gene circuit into a well-defined and well-expressed chromosomal locus, we utilized a genomic integration platform we recently established in HEK293 cells^12^. This platform overcomes major drawbacks of widely used viral and CRISPR-Cas9 based genomic integration methods by utilizing a Landing Pad (LP), which contains a set of selection markers and the GFP reporter flanked by a pair of heterologous Flp-recombinase targeting sites (FRT). We incorporated the LP into the AAVS1 safe harbor site (SHS)^30^ of the HEK293 human genome using CRISPR-Cas9-mediated target insertion **(Figure S1A)**. Once established, any desired DNA payload can be site-specifically integrated into the AAVS1-localized LP through Flp-recombinase-mediated cassette exchange (FLP-RMCE), without any mutations or other side effects. All of the gene circuits described henceforth **(Table S1)** were flipped into the same HEK293 LP parental clone, into a verified single LP copy in the AAVS1 site.

Using FLP-RMCE in HEK293-LP cells, we replaced the original LP cassette with the mNF-Cas13d gene circuit and new selection markers **(Figure S1B)**. Fluorescence shifting from green (GFP) to red (mCherry) in the surviving cell population indicated successful cassette switching via RMCE **(Figure S1B)**, with some basal expression even without doxycycline (Dox) induction **(Figure S1B)**.

To evaluate the Cas13d expression-tuning capabilities of the integrated mNF-Cas13d gene circuit **(Figure 1B)**, we monitored the expression mean and variability (coefficient of variation, CV) of mCherry reporter expression under constant Dox concentrations ranging from 0 to 100 ng/ml, 72 hours post-induction. We observed a gradual, monotonic, uniform increase in mCherry expression as indicated by narrow flow cytometry histograms **(Figure 1C)**. The Dox dose-response of mean mCherry expression indicated a 14-fold increase between maximum and basal levels **(Figure 1D)**. Additionally, the CV of mCherry expression remained consistently low, with a moderate increase only at high Dox concentrations **(Figure 1E)**, indicating stable, precise, low-noise characteristics as previously reported in other mNF gene circuits tuning the expression of other genes^12, 27, 31–32^.

### The mNF-Cas13d gene circuit dose-dependently reduces target mRNA, but with substantial basal effect

To rigorously assess the efficacy, tunabilty and precision of Dox dose-dependent target RNA reduction by the integrated mNF-Cas13d gene circuit, we co-expressed a BFP reporter with either non-targeting crRNA or BFP-targeting crRNA from a single-plasmid **(Figure S2A)**. Additionally, to determine the maximum possible target reduction, we engineered and integrated a mNF-disabled, constitutive Cas13d expression cascade identical to mNF-Cas13d except for lacking the TetO2 sites in the CMV promoter **(Figure S2A)**. We transfected both cell lines pre-induced or uninduced (Dox; 100 or 0 ng/ml for 72 hours) with plasmids containing either BFP-targeting or non-targeting crRNA along with the BFP reporter. Flow cytometry histograms indicated that only the mNF-Cas13d gene circuit could tune down RNA expression, reaching at full induction knockdown levels comparable to those of its constitutive counterpart **(Figure S2B)**. However, we observed a significant (>60%) unwanted basal target RNA reduction (henceforth, “basal effect”) even without induction at 0 ng/ml Dox, raising concerns about the applicability of the gene circuit.

To mitigate this pronounced basal effect, we asked if lower RNA target expression from the weaker SV40 promoter rather than the strong EF1α promoter could be helpful **(Figure 2A)**. The dose-dependent reduction in BFP levels indicated that, indeed, this new plasmid lowered the basal effect, but it still remained substantial (>40%, **Figure 2B-C)**, significantly limiting the dynamic range of BFP downregulation (ranging from ∼50% to ∼20% relative to control).

**Figure 2.**
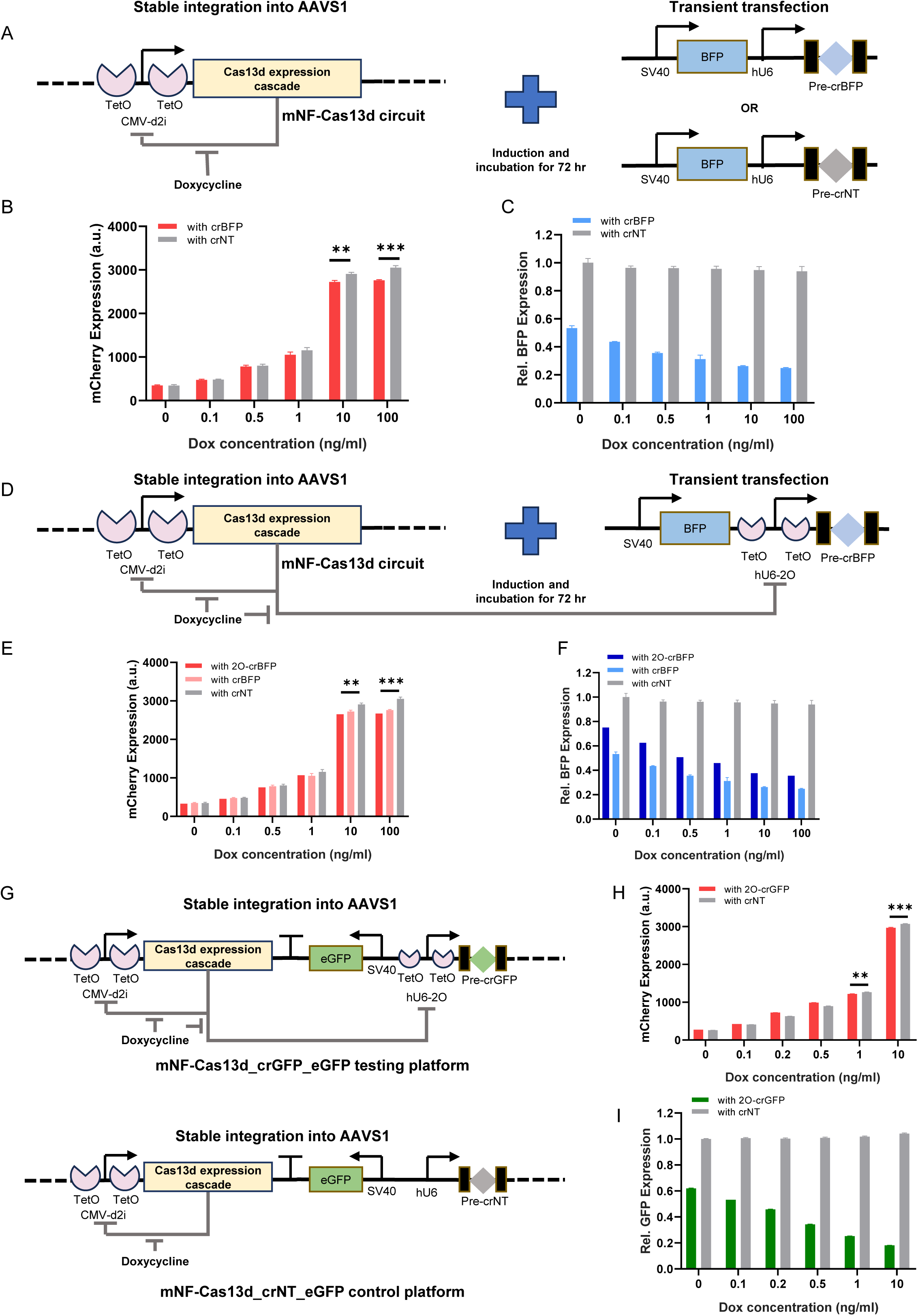
Negative-feedback transcriptional regulation of both Cas13d protein and crRNA improves dose-response characteristics. **(A)** Diagram of single-vector transfected to test Cas13d’s Dox dose-dependent on-target activity. The plasmid encoding the BFP reporter and the BFP-targeting crRNA or non-targeting crRNA was transfected into HEK 293 cells with mNF-Cas13d stably LP-integrated. Cells were Dox-induced and incubated for 72 hours before flow cytometry measurements. **(B)** Dox dose-responses of mean mCherry fluorescence intensity in HEK 293 cells with stably LP-integrated mNF-Cas13d gene circuit 72 hours post-transfection. Unpaired two-tailed t-test, n=3, **P<0.01, ***P<0.001. **(C)** Dox dose-dependent reduction of transiently transfected BFP reporter by Cas13d targeting. All BFP expression levels were normalized to uninduced cells with non-targeting guide, n = 3. **(D)** Diagram of single-vector transfection to test Cas13d’s Dox dose-dependent, on-target activity with regulated crRNA expression. BFP-targeting crRNA was driven by a modified human U6 (hU6) promoter containing two Tetracycline Operator (TetO2) sites flanking the TATA-box. This hU6-2O promoter can then be repressed by hTetR expressed from the mNF-Cas13d gene circuit. **(E)** Comparison of mCherry dose-responses from the mNF-Cas13d gene circuit 72 hours post-transfection with all three transfection constructs. Unpaired two-tailed t-test, n=3, **P<0.01, ***P<0.001. **(F)** Comparison of Dox dose-dependent BFP reduction by Cas13d, 72 hours post-transfection with all three transfection constructs. All BFP expression levels were normalized to uninduced samples with non-targeting guide, n = 3. **(G)** Diagram of all-in-one constructs to test Cas13d’s on-target activity on stably genome-integrated targets. The GFP reporter-targeting crRNA is expressed from the TetR-regulated hU6-2O promoter, while the non-targeting guide is freely expressed from the normal hU6 promoter. The whole construct was FLP-RMCE integrated using the same HEK 293 LP parental cells. **(H)** Dose-responses of mean mCherry fluorescence intensity for stably integrated all-in-one constructs in engineered HEK 293 cells 72 hours post-induction. Unpaired two-tailed t-test, n=3, **P<0.01, ***P<0.001. **(I)** Dose-dependent reduction of stably integrated GFP reporter expression by Cas13d. All GFP expression levels were normalized to uninduced samples with non-targeting guide, n = 3.

To further address this basal effect issue, we considered regulating crRNA levels, as opposed to the typical constitutive crRNA expression. We replaced the original constitutive hU6 promoter with a modified hU6-2O incorporating hTetR-repressible TetO2 sites^33^ (hU6-2O). Indeed, the basal effect significantly diminished in both mNF-Cas13d and constitutive Cas13d cells as indicated by the reduction of the strong EF1α promoter-driven BFP target expression (from over 60% to about 40%, **Figure S2C)**, while the fully induced, maximal reduction decreased only moderately from over 80% to about 75% **(Figure S2D)**. Furthermore, lower BFP target expression from the SV40 promoter also diminished the basal effect (from over 40% to about 25% relative to control) while increasing back the dynamic range of RNA downregulation (ranging from 75% to 35% relative to control) **(Figure 2D-F)**.

So far, we expressed the RNA targets from plasmids. To test the reduction of RNAs transcribed from the genome, we constructed and LP-integrated an all-in-one testing platform containing (i) the mNF-Cas13d gene circuit; (ii) a target GFP expressed from the SV40 promoter; and (iii) GFP-targeting crRNA expressed from the hU6-2O promoter. A similar system with a non-targeting crRNA served as control **(Figure 2G)**. We observed precise mCherry dose-responses from both integrated constructs **(Figure 2H)**, and a Dox dose-dependent GFP downregulation up to 80%, although a 40% basal effect persisted **(Figure 2I)**.

Overall, we found that the mNF-Cas13d gene circuit can reduce target RNA levels as expected, in a dose-dependent manner. However, we also observed a severe basal effect even without induction, related to basal expression of both Cas13d and crRNA, indicating the need of further optimization for broader applicability.

### The activated Cas13d:crRNA complex causes severe collateral off-target reduction

In all experiments so far, high induction led to significantly lower mCherry expression when accompanied with BFP-targeting crRNA, as compared to non-targeting crRNA **(Figure 2B, E, H)**. We hypothesized that collateral, off-target activity of the target-activated Cas13d:crRNA complex might be cutting mCherry, as observed previously for overexpressed targets, highly repetitive targets, and even sufficiently abundant endogenous genes^19, 29, 34^, with variations potentially dependent on cell type^20^ but evidenced in both HeLa and HEK293 cells^29^. Therefore, we set out to evaluate the extent of collateral RNA-cutting activity stemming from the mNF-Cas13d gene circuit, which might also contribute to the unwanted basal effect.

To examine the origins of collateral activity, we cloned an hTetR-regulated GFP-targeting crRNA together with a weaker SV40 promoter-driven non-target BFP reporter and the strong CMV promoter-driven GFP target, to boost up activated Cas13d:crRNA complexes **(Figure S3A)**. Upon co-transfection into mNF-Cas13d cells, GFP-targeting crRNA reduced the GFP target expression dose-dependently at both 1 and 10 ng/ml Dox induction **(Figure S3B-C)**, with a substantial basal effect **(Figure S3C)**, but mCherry and BFP expression also decreased consistently, indicating Cas13d’s collateral RNase activity **(Figure S3B,D)**.

In actual biological scenarios, the abundance of a specific target RNA is only a fraction of the total intracellular RNA pool. Therefore, to further assess collateral activity with systems resembling such scenarios, we modified the reporting system to express the non-target BFP reporter under the strong EF1α promoter, while expressing the GFP target from the weaker SV40 promoter **(Figure 3A)**. While dose-dependent GFP on-target reduction still correlated with increasing mCherry levels as intended **(Figure 3B-C)**, the dose-dependent reduction in BFP was more pronounced, and exacerbated by unregulated crRNA expression **(Figure 3D)**. This suggests that excessive activated Cas13d:crRNA complexes can cause significant off-target effects due to Cas13d’s collateral activity.

**Figure 3.**
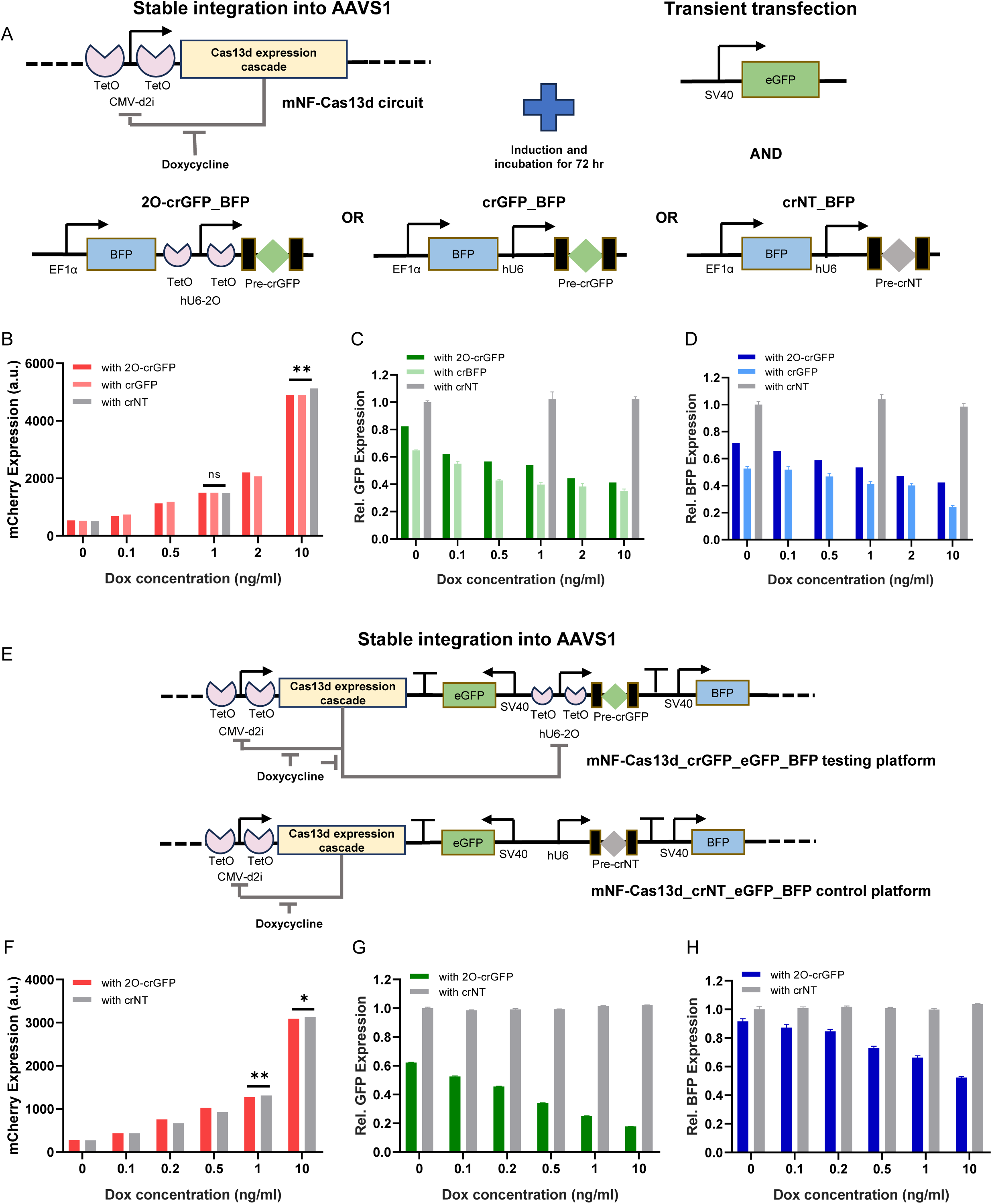
Collateral activity due to Cas13d hyperactivation depends on Cas13d and crRNA expression levels. **(A)** Diagram of experimental setup to assess collateral activity from hyperactivated Cas13d with different guide control scenarios. The SV40 promoter-driven GFP target was co-transfected with either TetR-regulated GFP-targeting crRNA, constitutively expressed GFP-targeting crRNA or non-targeting crRNA. All three crRNA plasmids contain the same BFP gene driven by the EF1α promoter. Cells were transfected under 72 hour induction before flow cytometry measurements. **(B)** Comparison of mCherry dose-responses from the mNF-Cas13d gene circuit 72 hours post-transfection for all three co-transfection scenarios. Unpaired two-tailed t-test, n=3, **P<0.01. **(C)** Comparison of relative GFP levels indicating dose-responses of on-target activity from the mNF-Cas13d gene circuit 72 hours post-transfection with all three co-transfection scenarios. All GFP expression levels were normalized to the uninduced sample with non-targeting guide, n=3. **(D)** Comparison of relative BFP levels indicating dose-responses of off-target activity from the mNF-Cas13d gene circuit 72 hours post-transfection with all three co-transfection setups. All the BFP expression levels were normalized to the uninduced sample with non-targeting guide, n=3. **(E)** Diagram of all-in-one constructs to test Cas13d’s on-target and off-target activities on stably genome-integrated targets. GFP reporter-targeting crRNA is expressed from the hTetR-regulated hU6-2O promoter, while the non-targeting guide is constitutively expressed from the normal hU6 promoter. The whole construct is FLP-RMCE-integrated using the same HEK 293 LP parental cells. **(F)** Dose-responses of mCherry reporter indicating Cas13d expression levels for stably integrated all-in-one constructs in HEK 293 cells 72 hours post-induction. Unpaired two-tailed t-test, n=3, *P<0.05, **P<0.01. **(G)** Dose-responses of relative GFP reporter levels indicating on-target activity for stably integrated all-in-one constructs in HEK 293 cells 72 hours post-induction. All GFP expression levels were normalized to uninduced samples with non-targeting guide, n=3. **(H)** Dose-responses of relative BFP reporter levels indicating off-target activity for stably integrated all-in-one constructs in HEK 293 cells 72 hours post-induction. All BFP expression levels were normalized to uninduced samples with non-targeting guide, n=3.

To map the fitness landscape of Cas13d system components, we evaluated how collateral activity affects cell viability. Using the reporter system with SV40-driven BFP and hTetR-regulated BFP-targeting crRNA **(Figure S3E)**, we found that activation of Cas13d:crRNA complexes is necessary for the observed effects as non-targeting crRNA did not significantly impact cell viability **(Figure S3F)**. However, on-target crRNA caused a Cas13d- and crRNA-dependent, dose-dependent decrease in cell viability, especially at higher induction **(Figure S3G)**, while the viability of cells transfected with non-targeting crRNA was practically unaffected.

Finally, to mimic a biological scenario where the mNF-Cas13d gene circuit reduces expression of specific gene targets neighboring non-target genes in the genome, we added an SV40-driven non-target BFP reporter to the all-in-one testing platform, with a non-targeting crRNA as control **(Figure 3E)**. The mCherry dose-responses from both target and control constructs had similar dynamic ranges with a drop in mCherry expression at higher induction **(Figure 3F)**. Dose-dependent GFP on-target downregulation reached up to 80%, with a 40% basal effect still persisting **(Figure 3G)**. There was a corresponding dose-dependent reduction in BFP levels, especially at higher induction, indicating collateral activity at substantial Cas13d and crRNA expression and activation **(Figure 3H)**.

Collectively, these results indicate substantial collateral activity when employing Cas13d to degrade sufficiently abundant RNA targets in cells, a challenge to be resolved before broad application in biological research and therapeutic development. The current mNF gene circuit design seems insufficient to address this issue, so further optimization is necessary.

### Multi-level negative autoregulation of both Cas13d and crRNA in MONARCH reduces collateral activity and improves on-target dynamic range

Given the excessive efficacy of both specific and nonspecific RNA degradation by the Cas13d:crRNA complex even at relatively low expression levels, we hypothesized that the key to reducing collateral activity and improving the dynamic range of on-target activity was to lower the expression ranges of both Cas13d and crRNA. Inspired by a previous study that utilized Cas13d’s crRNA processing capability as a mechanism for RNA-level negative autoregulation^29^, we incorporated a pre-crRNA motif into the 3’UTR of the mNF coding region before the poly(A) signal. This innovation led to the design of a multi-level optimized negative-autoregulated Cas13d and crRNA hybrid circuit, termed MONARCH **(Figure 4A)**.

**Figure 4.**
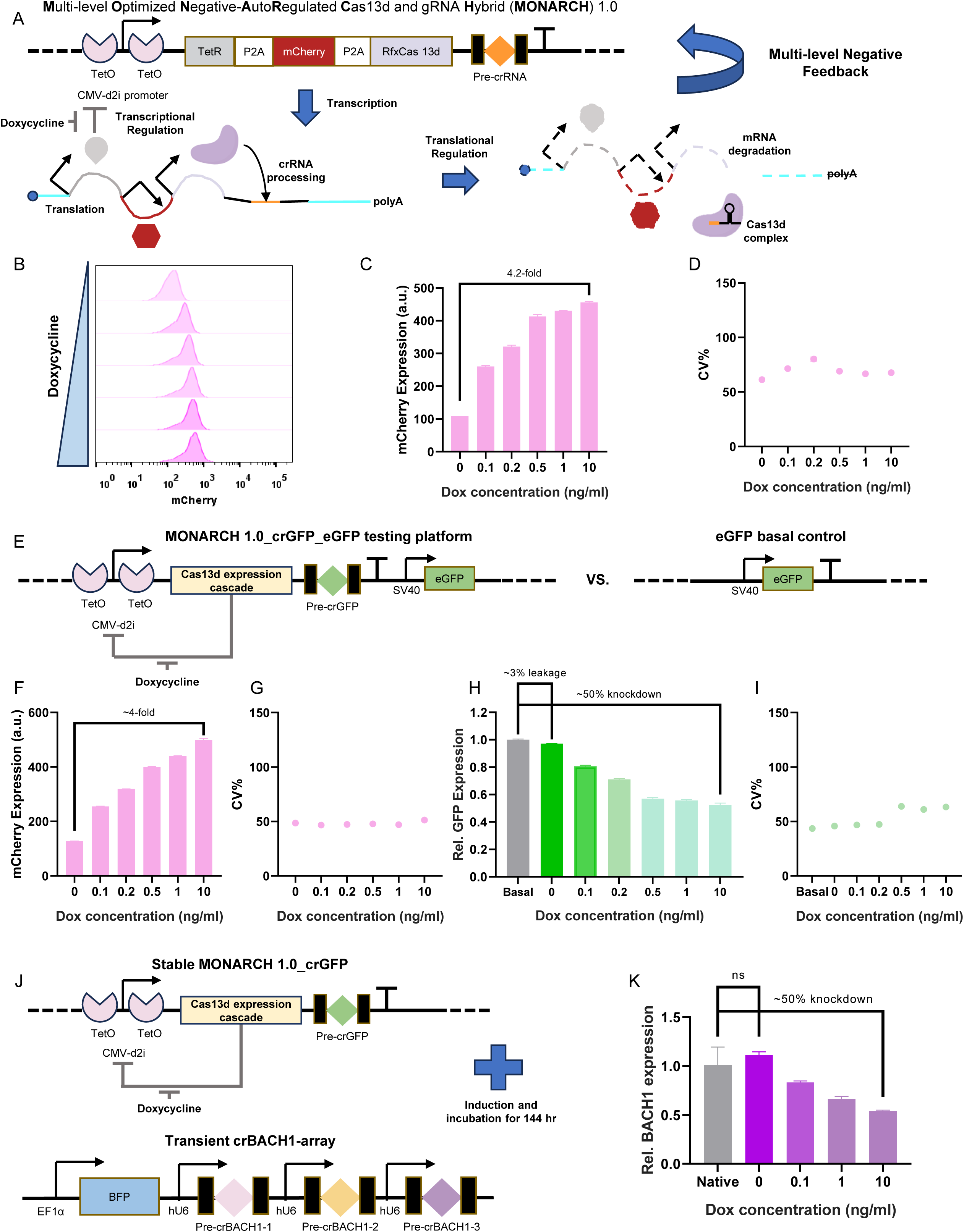
Multi-level negative autoregulation of Cas13d and crRNA in MONARCH reduces the basal target downregulation. **(A)** Design and rationale for multi-level negative autoregulation of both Cas13d and crRNA expression, to avoid their unwanted overexpression, geberating too many activated Cas13d:crRNA complexes with excessive basal effect and collateral activity. Incorporating the crRNA into the same transcript with Cas13d not only brings both under the same tight transcriptional regulation, but also adds RNA-level regulation via crRNA processing by Cas13d. This may reduce the basal effect as well as the collateral activity. **(B)** Representative dose-responses of fluorescence intensity histograms from the stably integrated MONARCH 1.0 circuit measured at 0, 0.1, 0.2, 0.5, 1, 10 ng/ml Dox levels, respectively. **(C)** Dose-responses of mean mCherry fluorescence intensity for the MONARCH 1.0 circuit stably integrated into HEK 293 LP cells (n=3). **(D)** Dose-responses of the coefficient of variation (CV) of mCherry reporter expression for MONARCH 1.0 stably integrated into HEK 293 LP cells (n=3). **(E)** Diagram for testing on-target activity of Cas13d expressed from MONARCH 1.0 on stably genome-integrated targets. The whole construct is FLP-RMCE-integrated using the same HEK 293 LP parental cells. Basal level is determined with integration of only the SV40 promoter-driven GFP target. **(F)** Dose-responses of mean fluorescence intensity of mCherry reporter for stably integrated MONARCH 1.0_SV40-GFP construct in HEK 293 cells (n=3). **(G)** Dose-responses of coefficient of variation (CV) of mCherry reporter for stably integrated MONARCH 1.0_SV40-GFP in HEK 293 cells (n=3). **(H)** Dose-responses of relative GFP levels indicating on-target activity for the MONARCH 1.0_SV40-GFP construct stably LP-integrated into HEK 293 cells. All GFP expression levels were normalized to the basal sample, n=3. **(I)** Dose-responses of coefficient of variation (CV) of GFP target expression for MONARCH 1.0_SV40-GFP stably LP-integrated into HEK 293 cells (n=3). **(J)** Diagram for testing on-target activity of MONARCH 1.0 on an endogenous target. GFP-targeting crRNA serves only as an RNA-level regulator in this scenario. Multiple crRNAs targeting BACH1 are cloned into a single plasmid and transfected together into the cells. RT-qPCR was performed after 144 hours of induction and incubation post-transfection. **(K)** Dose-dependent reduction of endogenous BACH1 by Cas13d expressed from MONARCH 1.0 stably LP-integrated into HEK 293 cells. One-way ANOVA with Tukey’s multiple comparisons tests between each dose and native control, n=3.

Like mNF, the MONARCH 1.0 system has transcriptional autorepression through hTetR binding to TetO2 sites at the promoter region, while Cas13d expression accelerates the degradation of its own mRNA, through poly(A) signal elimination when processing the crRNA within the 3’UTR. This design lowers both Cas13d and crRNA expression levels at each induction dosage and thus reduces the basal effect **(Figure 4A)**. To characterize the performance of this new gene circuit, we integrated MONARCH 1.0 into the AAVS1 site of HEK 293 LP parental cells using the same RMCE strategy. Upon Dox induction, mCherry levels increased in a Dox dose-responsive manner as evidenced by histogram shifts and fluorescence quantitation **(Figures 4B-C)**, while preserving a precise, low-noise expression profile characteristic of mNF gene circuits **(Figure 4D)**. The total tunability range of Cas13d expression shifted down compared to traditional mNF circuits, as expected due to stronger RNA-level destabilization by poly(A) loss at higher dosages **(Figure 4C)**.

To determine whether collateral activity still persists in the MONARCH 1.0 system, we co-transfected the cells with an abundant GFP target and a moderate non-target BFP reporter **(Figure S4A)**. Compared to the mNF-Cas13d gene circuit with non-targeting crRNA control, the levels of mCherry expression dropped significantly **(Figure S4B)**, lowering the basal effect further than for the regulated crRNA strategy (20% vs. 30% relative to control). GFP expression dropped as expected **(Figure S4C)**, while BFP expression stayed practically unchanged, indicating vanishing collateral activity **(Figure S4D)**. Overall, MONARCH 1.0 successfully reduces unwanted collateral activity while preserving dose-dependent on-target activity with even lower basal effect.

Furthermore, to assess the Dox dose-dependent downregulation of on-target and off-target RNA expressed from LP-integrated targets with MONARCH 1.0, we AAVS1-inserted all-in-one MONARCH 1.0 platforms **(Figure 4E and Figure S4E)**. MONARCH downshifted the dynamic range of mCherry expression again, indicating Cas13d and crRNA level reduction **(Figures 4F and Figure S4F)**, with low-noise throughout **(Figures 4G and Figure S4G)**. The basal effect dropped to only 3-5%, while the maximum on-target reduction remained about 50% from the basal **(Figures 4H and Figure S4H)**. GFP-target expression variation remained low, suggesting homogeneous RNA degradation by Cas13d across the cell population **(Figure 4I)**. Additionally, we observed no detectable collateral activity on the non-target BFP reporter while reducing GFP dose-dependently **(Figure S4I)**.

Finally, after demonstrating improved on- and off-target downregulation characteristics with the MONARCH 1.0 system, we tested it on an actual endogenous gene target, BACH1, an important metastatic regulator in breast cancer cells^28,34^. By combining multiple crRNAs and transiently transfecting them into MONARCH 1.0 cells **(Figure 4J)**, we observed a Dox-dependent, gradual reduction in BACH1 mRNA (up to 50% from the basal) without a basal effect **(Figure 4K)**. This demonstrated the MONARCH system’s potential to achieve precise and effective native target RNA downregulation, even transiently.

Overall, we have introduced a novel gene circuit design that leverages multi-level negative autoregulation on Cas13d and crRNA to simultaneously reduce collateral activity and improve on-target activity characteristics. The MONARCH system’s capability to tune down both transient and stable RNA targets will offer significant potential for quantitative genetic studies and therapeutic development.

### MONARCH’s optimization improves its on-target activity while preserving minimal collateral activity

We have demonstrated that MONARCH 1.0 retains dose-dependent target RNA downregulation without detectable collateral activity. However, we considered the maximum target reduction of ∼50% unsatisfying, which could worsen for lower-expressed native targets. Therefore, we sought to improve MONARCH’s on-target activity, especially at higher induction levels, without reintroducing collateral activity. We hypothesized that Cas13d and crRNA expression may have become too low in MONARCH 1.0, so elevating them slightly could improve on-target activity without significant collateral activity. Thus, we designed MONARCH 2.0 **(Figure 5A)**.

**Figure 5.**
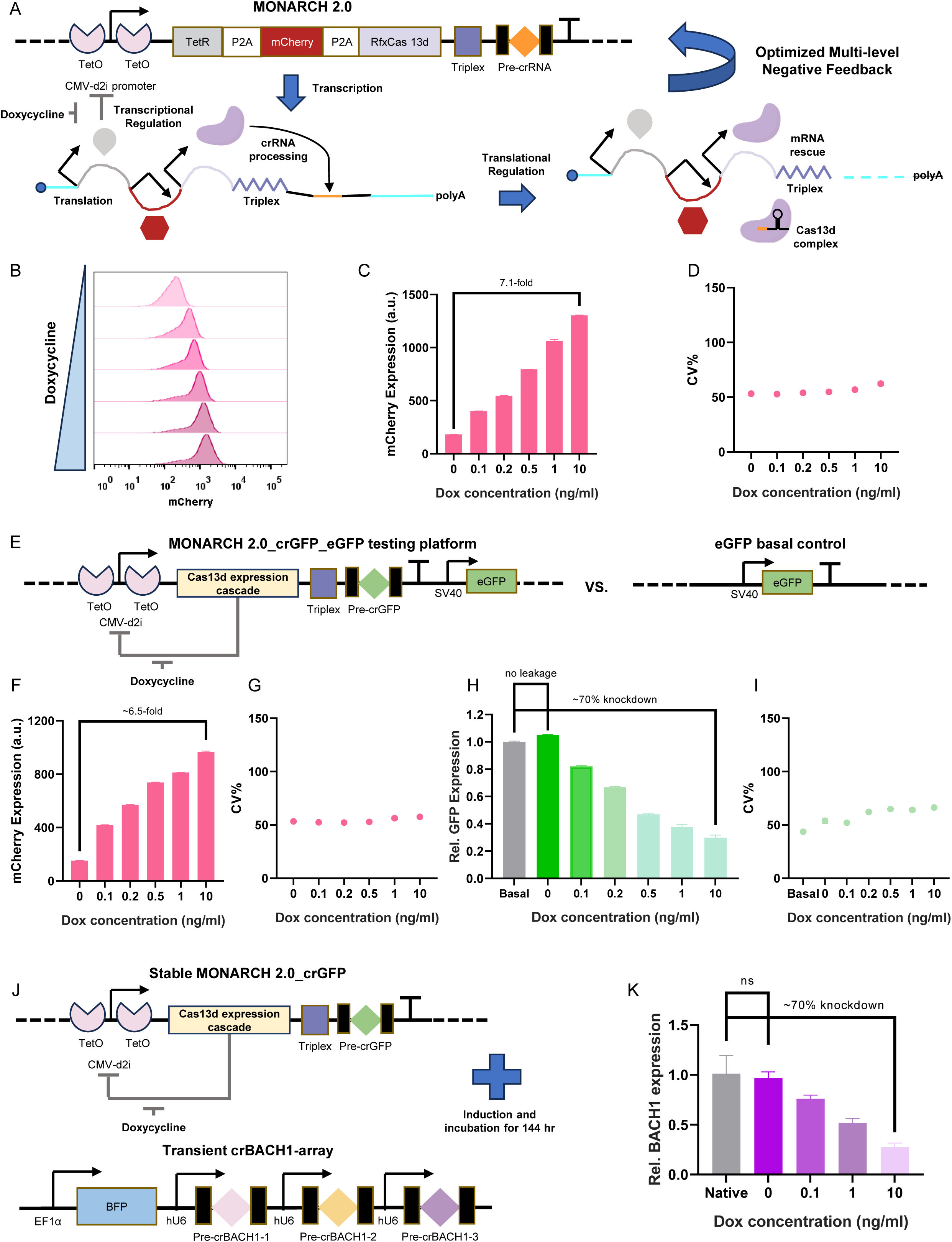
Optimized multi-level negative autoregulation of Cas13d and crRNA improves the on-target potency and minimizes the basal effect. **(A)** Design and rationale for optimized multi-level negative-autoregulation for Cas13d and crRNA expression. Compared to MONARCH 1.0, a tertiary RNA structural motif “Triplex” is incorporated between the CDS and crRNA to stabilize transcripts lacking poly(A) due to crRNA processing, enabling a moderately extended window of protein expression. **(B)** Representative dose-responses of fluorescence intensity histograms from the stably integrated MONARCH 2.0 circuit measured at 0, 0.1, 0.2, 0.5, 1, 10 ng/ml Dox levels, respectively. **(C)** Dose-responses of mean mCherry fluorescence intensity for the MONARCH 2.0 circuit stably LP-integrated into HEK 293 cells (n=3). **(D)** Dose-responses of coefficient of variation (CV) of the mCherry reporter for MONARCH 2.0 stably LP-integrated into HEK 293 cells (n=3). **(E)** Diagram for testing the on-target activity of Cas13d expressed from MONARCH 2.0 on genomically integrated targets. The whole construct is FLP-RMCE-integrated using the same HEK 293 LP parental cells. The basal level is determined through the integration of only SV40 promoter-driven GFP. **(F)** Dose-responses of mean mCherry fluorescence intensity for the MONARCH 2.0_SV40-GFP construct stably LP-integrated into HEK 293 cells (n=3). **(G)** Dose-responses of coefficient of variation (CV) of the mCherry reporter for MONARCH 2.0_SV40-GFP stably LP-integrated into HEK 293 cells (n=3). **(H)** Dose-responses of relative GFP levels indicating on-target activity for MONARCH 2.0_SV40-GFP stably LP-integrated into HEK 293 cells. All the GFP expression levels were normalized to the basal sample, n=3. **(I)** Dose-responses of coefficient of variation (CV) of the GFP target for stably integrated MONARCH 2.0_SV40-GFP in HEK 293 cells (n=3). **(J)** Diagram for testing on-target activity of MONARCH 2.0 on an endogenous target. GFP-targeting crRNA serves only as an RNA-level regulator in this scenario. Multiple crRNAs targeting BACH1 are cloned into a single plasmid and transfected together into the cells. RT-qPCR was performed after 144 hours of induction and incubation post-transfection. **(K)** Dose-dependent reduction of endogenous BACH1 by Cas13d expressed from MONARCH 2.0 stably LP-integrated into HEK 293 cells. One-way ANOVA with Tukey’s multiple comparisons test between each dose and native control, n=3.

MONARCH 2.0 was almost identical to MONARCH 1.0, except we incorporated Triplex, an additional RNA structural motif between the coding region and pre-crRNA motif. Triplex is a 110-nt structure derived from the 3′ end of the mouse non-coding RNA Metastasis-associated lung adenocarcinoma transcript 1 (Malat1), previously found to stabilize transcripts lacking poly(A) tails through the formation of a tertiary structure^35^. It has been successfully used to incorporate CRISPR-Cas12a and its guide RNA into a single transcript^36^. Here, we similarly applied it to enable better Cas13d expression with crRNA in the same transcript **(Figure 5A)**.

To characterize the performance of MONARCH 2.0, we integrated MONARCH 2.0 into the AAVS1 site in HEK 293 LP parental cells using the same RMCE strategy. Upon Dox induction, histogram shifts and fluorescence quantitation indicated significantly improved dose-responsive tuning of mCherry levels **(Figures 5B-C)**, while preserving a low-noise expression profile throughout **(Figure 5D)**. With MONARCH 2.0, Cas13d expression was tunable over a 7-fold dynamic range, with minimal basal expression comparable to that of MONARCH 1.0, but significantly higher maximal Cas13d expression at full induction **(Figure 5C)**.

To determine whether MONARCH 2.0 would reintroduce collateral activity, we again co-transfected the cells with an abundant GFP target and a moderate non-target BFP reporter **(Figure S5A)**. First, compared to the mNF-Cas13d gene circuit with non-targeting crRNA control, we still observed significantly lower levels of mCherry expression **(Figure S5B)**, and basal effect as low as for MONARCH 1.0. Dose-dependent GFP reduction improved **(Figure S5C)**, without collateral activity except for a slight increase (5% change in BFP expression) only at the highest induction dosage **(Figure S5D)**, which did not compromise cell viability **(Figure S3G).** Considering that the GFP target is overwhelmingly abundant, native targets should be safe from collateral cutting. Overall, the MONARCH 2.0 system effectively improved on-target activity characteristics while still preserving minimal collateral activity.

To assess the safety and effectiveness of MONARCH 2.0 in a close-to-real scenario, we stably integrated a target gene along with stably integrated all-in-one MONARCH 2.0 platforms **(Figure 5E and Figure S5E)**. The dynamic range of mCherry expression improved, indicating partially restored levels of Cas13d and crRNA at higher dosages **(Figures 5F and Figure S5F)**. Noise was low throughout **(Figures 5G and Figure S5G)**, and the basal effect diminished further, to non-detectable levels. We hypothesized that partially restored mRNA stability provided more TetR protein, causing stronger transcriptional repression of crRNA expression, especially without induction **(Figure 5A)**. The maximum on-target reduction reached up to 70% from the basal, approaching mNF-Cas13d’s 80% **(Figures 5H and Figure S5H)**. Similarly, GFP-target expression variation remained stably low, suggesting homogeneous RNA degradation across the cell population **(Figure 5I)**. More importantly, there was no detectable collateral activity on the non-target BFP reporter, suggesting that MONARCH 2.0 remains safe for downregulation of most RNA targets **(Figure S5I)**.

Finally, we tested how the action of MONARCH 2.0 on the same endogenous gene target BACH1, compared to MONARCH 1.0. By transiently transfecting the same amount of BACH1 crRNA array into MONARCH 2.0 cells **(Figure 5J)**, there was no basal effect from uninduced MONARCH 2.0, while tunable BACH1 mRNA reduction improved up to 70% from the basal **(Figure 5K)**, consistent with observations from reporter RNA targeting. This demonstrates consistent improvements of the MONARCH 2.0 system over MONARCH 1.0, useful for achieving more effective and precise target RNA downregulation.

Overall, by incorporating Triplex, an RNA stabilizing element we improved the on-target activity of MONARCH, generating the upgraded MONARCH 2.0 system. We demonstrated that MONARCH 2.0 effectively improved maximal target reduction while preserving minimal collateral activity, indicating a strong potential for broad applications in genetic studies and therapeutic developments.

### Mathematical model captures MONARCH’s benefits and indicates target RNA cutting by nonspecific Cas13d activity

So far, MONARCH has had three main benefits: it eliminated the basal effect and collateral cutting while improving the dynamic range of target reduction. To explain the improved performance of the MONARCH system, we hypothesized that Cas13d-mediated nonspecific RNA cutting significantly reduces the target RNA, too. Further, we assumed that nonspecific cutting is primarily directed towards target RNA at low Cas13d-crRNA complex levels, whereas an excess of such activated complexes shifts nonspecific cutting towards mostly non-target RNA **(Figure 6A)**. Thus, lowering Cas13d-crRNA amounts restored on-target efficacy despite significantly lower Cas13d and crRNA expression, thereby minimizing the collateral activity. Based on protein-conformational data^37^, we will refer to the target-free Cas13d-crRNA duplex as “active”, and to the RNA-target-bound Cas13d-crRNA triplex as “hyperactive”.

**Figure 6.**
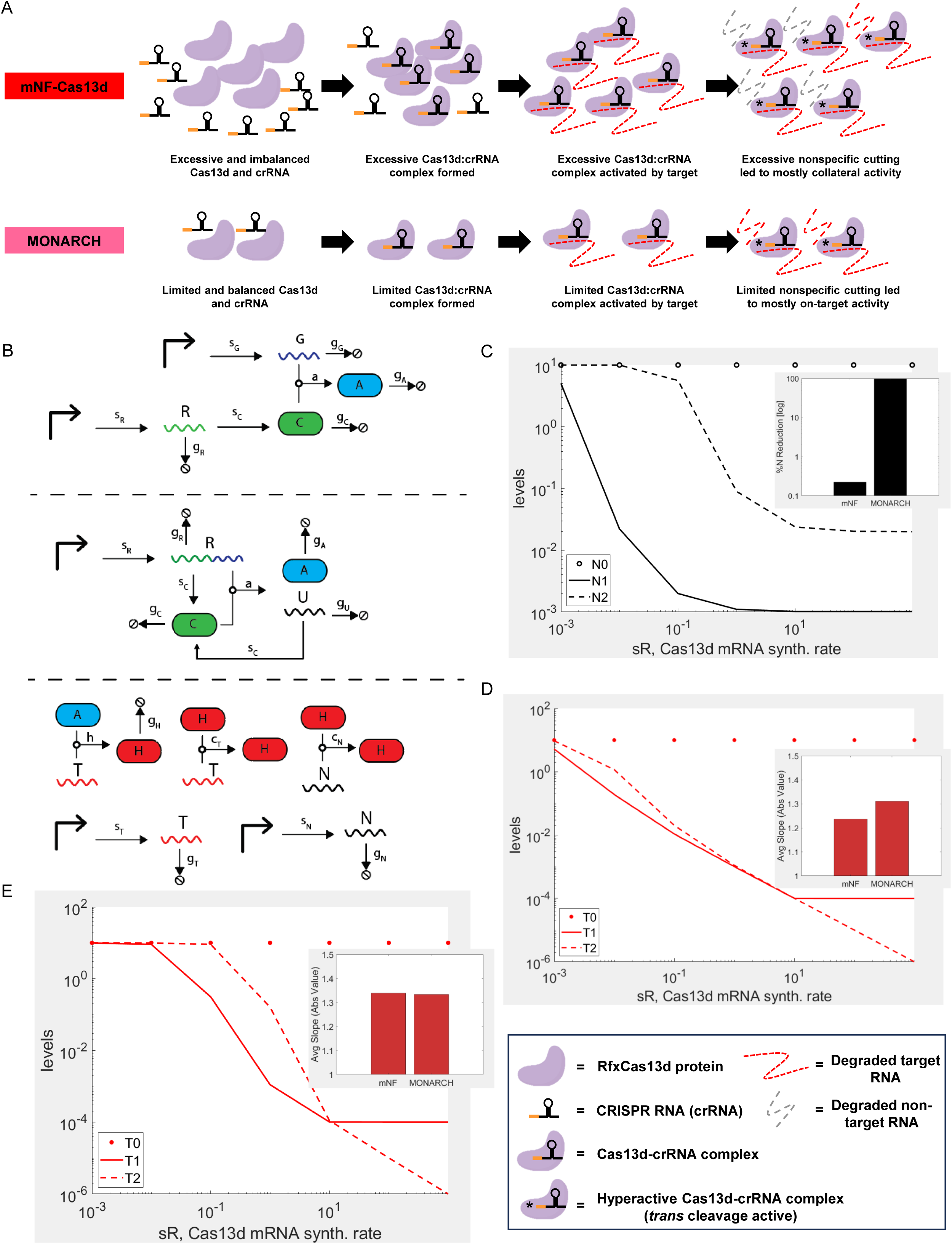
Mathematical modeling indicates the origins of MONARCH’s benefits. (A) Illustration of hypothesized mechanisms to explain experimental observations from both mNF-Cas13d and MONARCH system. Briefly, with mNF-Cas13d, highly expressed Cas13d and crRNA will form excessive amounts of Cas13d:crRNA complexes, which can be hyperactivated by target RNA cutting. Due to the limited target RNA pool, the excessive hyperactive Cas13d:crRNA complexes will start to degrade non-target RNA, causing collateral activity. With MONARCH, the Cas13d:crRNA complex abundance is limited by multi-level negative autoregulation and remains sufficiently curtailed compared to the target RNA pool. Then the limited amount of hyperactive complex will mostly cut target RNA, resulting in an elevated on-target efficacy while minimize the collateral activity. Red dashed curve, degraded target RNA; grey dash curve, degraded non-target RNA; “*” indicates nonspecific, *trans* cleavage activation (complex hyperactivation). (B) Schematic of modeling reactions. *Top*: reactions to produce the active Cas13d-crRNA complex (*A*) in the mNF-Cas13d system. Here, the guide RNA (*G*) is produced from a separate promoter from the Cas13d transcript (*R*). The Cas13d transcript is translated into Cas13d protein (*C*), which then associates with the guide RNA to become activated. *Middle*: reactions to produce the active Cas13d complex in the MONARCH system. Here, the guide RNA is co-expressed with the Cas13d transcript. When the Cas13d protein processes the RNA, it produces an unstable Cas13d transcript (*U*), which can still be translated into Cas13d protein. *Bottom*: for both models, once the active Cas13d complex is created, it can then start cutting target RNA (*T*). When this happens, the Cas13d complex becomes hyperactive (*H*), allowing it to cleave both target RNA and nonspecific RNA (*N*). (See **Supplemental Results** for more details). (C) Plot of simulated nonspecific RNA levels for each system. *N0* (circles): no Cas13d control (i.e. only constitutive synthesis and linear degradation). *N1* (full line): mNF-Cas13d system. *N2* (dashed line): MONARCH system. Bar chart shows the relative basal levels, calculated relative to *N0*. (D) Plot of simulated target RNA levels for each system. *T0* (circles): no Cas13d control (i.e. only constitutive synthesis and linear degradation). T*1* (full line): mNF-Cas13d system. *T2* (dashed line): MONARCH system. Bar chart shows the average slope, calculated from the dynamic range of each system (See the **Supplemental Results** for more details). (E) Repeat simulation with cT (cutting rate of target RNA by hyperactive Cas13d) set to zero, indicating that nonspecific RNA cutting contributes to target RNa reduction. Legend is the same as in **(D)**.

To test this hypothesis, we established mathematical models based on mass action kinetics for both the mNF-Cas13d and MONARCH systems (See **Supplemental Results** for detailed descriptions). After defining and parametrizing the underlying biochemical reactions **(Figure 6B)**, we studied the effect of increasing Dox levels by increasing the Cas13d mRNA synthesis rate over a range of several orders of magnitude. Indeed, we observed a significant decrease of collateral activity by nonspecific RNA cutting in MONARCH compared to mNF-Cas13d models **(Figure 6C)**. Next, we also found a decrease in the basal effect of MONARCH compared to mNF-Cas13d, while MONARCH still reached similar maximal on-target efficacy **(Figure 6D)**. To quantify this improved dynamic range, we calculated the average slope for target RNA cutting versus Cas13d mRNA synthesis rate. Indeed, MONARCH had a higher slope, indicating its wider dynamic range **(Figure 6D)**. These simulation results corresponded well with the experimentally observed benefits of MONARCH **(Figure S6A).** Interestingly, this improved dynamic range was only captured when we assumed that nonspecific RNA cutting also degraded target RNA, suggesting such improvement was due to target RNA-cutting by hyperactive Cas13d **(Figure 6E)**.

This latter observation led to the prediction that removing Cas13d’s nonspecific RNA cutting could compromise its overall on-target efficacy **(Figure S6B-C)**. To test this prediction experimentally, we co-transfected a plasmid that constitutively expressed collateral activity-free, high-fidelity RfxCas13d (hfCas13d)^34^, and individual crRNA together with a EF1α-driven BFP reporter into HEK293 cells **(Figure S6B)**. We observed no detectable mCherry expression changes with all three crRNAs **(Figure S6C)**, and eGFP or BFP expression only dropped upon introducing the corresponding crRNA **(Figure S6D-E)**, indicating an absence of collateral activity. However, compared to constitutively expressed wildtype RfxCas13d, hfCas13d had significantly lower maximal target RNA reduction (80% vs. 50% relative to control) **(Figure S2B and Figure S6D)**. Overall, these results corroborated that nonspecific activity of hyperactive Cas13d beneficially contributed to its target RNA reduction efficacy.

### MONARCH demonstrates potential for tunable anti-viral activity in engineered Vero E6 cells

Cas13d, an RNA-guided RNA-targeting nuclease without protospacer flanking sequence (PFS) constraints^38^, has significant potential as an artificial antiviral effector^39^. However, both our research and other studies have identified that unregulated overexpression of Cas13d and crRNAs can cause substantial collateral activity, severely compromising its utility in therapeutic applications and biological research. Therefore, appropriate regulation of Cas13d is crucial for its safe deployment in any intended application. Here, we illustrate the potential of the MONARCH system as a regulatable, side effect-free antiviral defense system against the COVID-19-causing SARS-CoV2 virus at the cellular level.

While direct experimentation with the full SARS-CoV-2 virus is challenging, we explored the potential of targeting viral genomic fragments using the MONARCH system engineered into Vero E6 green monkey cells, a standard model for human viral studies. We first assessed the feasibility of employing the same integration techniques used in human cells for generating a landing pad platform at the AAVS1 site of Vero E6 cells. Intriguingly, BLAST alignment revealed a corresponding AAVS1 ortholog in the African Green Monkey (*Chlorocebus aethiops*) genome, from which the Vero E6 cell line was derived, with a preserved PAM site for the sgRNA used in human cells, albeit with two distant mismatches **(Figure S7A)**. This alignment suggested the potential use of the same LP generation pipeline and sgRNA sequences for LP insertion into the AAVS1 ortholog in Vero E6 cells.

We proceeded as before, using enhanced SpCas9 (eSpCas9) to integrate the LP cassette into the Vero E6 genome, specifically at the AAVS1 ortholog site **(Figure S7B)**. We used an RFP reporter cloned into the LP donor vector backbone to negatively select against random integration. Following puromycin selection, we successfully isolated a group of surviving Vero cells, all expressing only green fluorescence, from which we established single-cell derived clonal populations **(Figure S7C)**. High and stable GFP expression in these clones suggested that the AAVS1 ortholog might also serve as a safe harbor site in the Vero genome. Subsequently, we employed the same RMCE strategy to site-specifically integrate two MONARCH systems co-expressing crRNA arrays targeting SARS-CoV-2 genome fragments, into the AAVS1 ortholog **(Figure 7A and Figure S7D)**.

**Figure 7.**
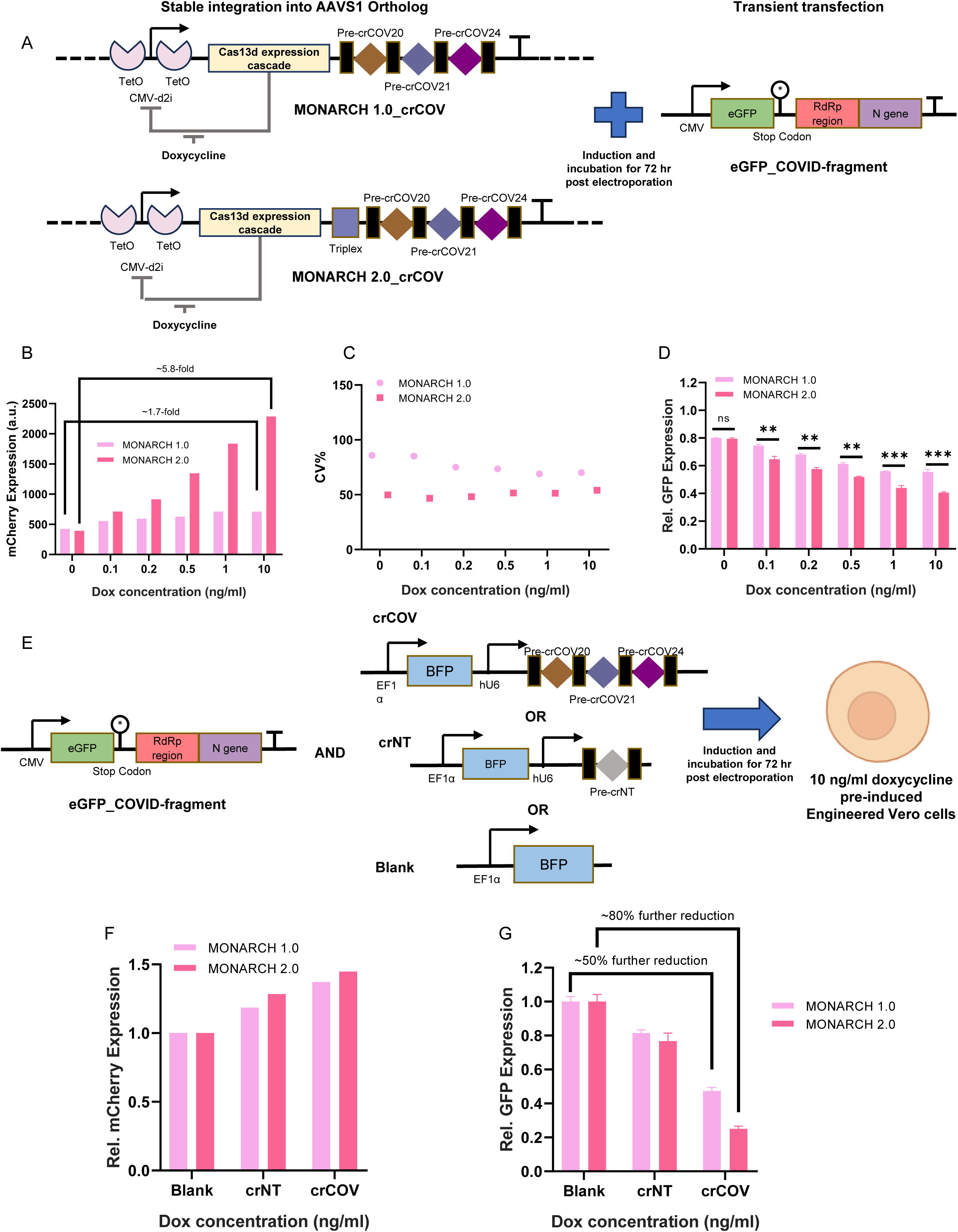
MONARCH established in Vero E6 LP cells demonstrates anti-viral capability and potential for improving on-target efficiency. **(A)** Diagram of experimental setup to repurpose MONARCH to target SARS-CoV-2 viral genome fragments in engineered Vero E6 cells. Both MONARCH systems are stably integrated into the AAVS1 ortholog in the Vero E6 genome using the same Landing-Pad and RMCE strategy. Successful by Cas13d targeting of SARS-CoV-2 viral genome fragments cloned into the 3’UTR of the GFP transcript will result in mRNA degradation, which can be measured by GFP fluorescence reduction. **(B)** Comparison of mCherry dose-responses from MONARCH 1.0 and 2.0 stably LP-integrated into Vero E6 cells 72 hours post-transfection. **(C)** Comparison of coefficient of variation (CV) of mCherry dose-responses from MONARCH 1.0 and 2.0 stably integrated in Vero E6 cells 72 hours post-transfection. **(D)** Comparison of relative GFP levels indicating dose-responses of on-target activity from MONARCH 1.0 and 2.0 stably LP-integrated into Vero E6 cells, 72 hours post-transfection. All the GFP expression levels were normalized to native Vero E6 cells transfected with the target plasmid only. Unpaired two-tailed t-test for each dosage comparison, n=3, **P<0.01, ***P<0.001. **(E)** Diagram of experimental setup to assess further improvement of MONARCH on-target activity by providing extra crRNA. Target donor plasmid is co-transfected with either SARS-CoV-2-targeting CRISPR array, non-targeting crRNA or blank control into the engineered Vero E6 cells. Engineered Vero E6 cells are pre-induced with 10 ng/ml doxycycline 72 hours before transfection. **(F)** Relative mCherry expression increases from 10 ng/ml doxycycline pre-induced MONARCH 1.0 and 2.0 stably integrated into Vero E6 cells, 72 hours post-transfection with extra crRNA plasmids. All mCherry expression levels were normalized to the Vero E6 cells transfected with a blank control plasmid. **(G)** Relative GFP level reductions 72 hours post-transfection with extra crRNA plasmids. All GFP expression levels were normalized to the Vero E6 cells transfected with a blank control plasmid.

Additionally, we constructed an all-in-one platform reporter system for SARS-CoV-2 genomic fragments, incorporating partial RdRp and N genes from the SARS-CoV-2 genome into the 3’UTR of a CMV-GFP expression cascade **(Figure 7A)**. Following transfection of Vero E6 cells engineered with both the MONARCH system and LP-free wildtype cells as controls, we observed that mCherry levels increased in a dose-responsive manner in both MONARCH systems, with comparable dynamic ranges to those of MONARCH 1.0 and 2.0 in human cells **(Figure 7B)**. Both circuits had low CV across all dosages **(Figure 7C)**. Furthermore, both systems demonstrated the capacity to reduce GFP reporter levels in a dose-dependent manner, indicative of an inducible reduction of viral fragments within the Vero E6 cells **(Figure 7D)**.

To further enhance the effectiveness of the MONARCH system, we hypothesized that providing additional crRNA arrays could amplify the reduction of viral fragments by introducing extra targeting crRNAs and creating a decoy effect to reduce Cas13d self-degradation. We tested this by co-transfecting the SARS-CoV-2 reporter cascade with either SARS-CoV-2-targeting crRNA arrays, non-targeting crRNA, or a blank control into 10 ng/ml Dox pre-induced, MONARCH-integrated Vero cells. The results confirmed our hypothesis, as even non-targeting crRNA induced more Cas13d expression due to the decoy effect, compared to the blank control, while the additional array of crRNAs caused stronger Cas13d induction **(Figure 7F)**. Notably, by transiently introducing extra targeting crRNAs into the cells, on-target activity was further elevated by about 50% and 80% for MONARCH 1.0 and 2.0 respectively, resulting in over 75% and 90% total reduction of viral fragments **(Figure 7G)**.

Overall, these findings suggest that MONARCH could be effectively leveraged as a regulatable antiviral defense system at the single-cell level. Additionally, the LP-engineered Vero E6 cell lines could support broader, quantitative viral studies with a reliable site for replaceable genomic integration, enabling precise, mutation-free, single-copy, site-specific DNA payload insertion and versatile repurposing.

## DISCUSSION

The discovery of Cas13d has sparked considerable interest due to its unique RNA-targeting capabilities, with broad potential for biotechnological and clinical applications. However, Cas13d’s nonspecific RNA-cutting *trans* activity, conformationally triggered upon binding its target RNA, has raised concerns due to collateral RNA degradation, causing global transcriptomic alterations and cell viability decline^29^ **(Figure S3G)**. Notably, the extent of collateral activity has been linked to target RNA abundance, being severe for highly overexpressed, repetitive targets, but negligible for low-expression targets^29^, in agreement with our data **(Figures 3C-D and Figure S3C-D)**. This link to target RNA abundance may explain the disparate findings for Cas13d’s collateral activity, ranging from negligible^40–42^ to severe^20, 29, 43^ in mammalian cells. Despite these issues, the potential utility of target RNA downregulation by Cas13d as a modular, programmable alternative to traditional RNA interference technologies should not be overlooked. Moreover, some studies have proposed harnessing Cas13d’s collateral activity for inducing cancer cell-specific apoptosis through global RNA depletion^44–45^. Therefore, it is crucial to assess the extent of nonspecific, *trans* cleavage and its separate contributions to on- and off-target cutting when employing Cas13d, to properly interpret its biological effects in various experimental contexts.

To address these questions, here we developed synthetic gene circuits that tune the levels of Cas13d and crRNA in various ways, with unprecedented sophistication. Our experimental and mathematical modeling data suggest that the expression levels of both Cas13d and crRNA are critical in modulating on-target and collateral activity. For example, even minimal amounts of the Cas13d:crRNA complex can act efficiently on its target RNA substrate **(Figure 4H and Figure S4C)**. Furthermore, both on-target and collateral activity strengthen for highly expressed Cas13d and crRNA **(Figure 3C-D, G-H and Figure S3C-D)**. Given that nonspecific, *trans* cleavage is an inherent characteristic of Cas13d triggered by binding its RNA target^37^, a mathematical model we developed indicated that *trans* cleavage may contribute critically to the efficacy of on-target degradation. Therefore, Cas13d’s *trans* cleavage (nonspecific RNA cutting) strongly degrades target RNAs, in addition to its *cis* cleavage (specific RNA cutting). Concordantly, the relative abundance of RNA target versus hyperactive Cas13d-crRNA-targetRNA triplex is crucial to minimize unwanted collateral activity, emphasizing the importance of synthetic biological expression tuning. In consensus with the recent finding of weaker on-target activity for high-fidelity RfxCas13d^34^ compared to the wildtype RfxCas13d^46^, we found that nonspecific RNA cutting, when properly controlled, can synergize with specific RNA cutting (on-target) activity to improve target RNA downregulation by Cas13d.

To leverage the contribution of nonspecific cutting to target RNA downregulation, we developed MONARCH 1.0, a multi-level negative-autoregulation circuit designed to finely tune the expression of both Cas13d and crRNA **(Figure 4A)**. Although effective at reducing both collateral activity and basal effect, the initial MONARCH excessively curtailed Cas13d and crRNA expression, inadvertently limiting maximal on-target activity **(Figures S4H-I)**. Seeking to balance expression levels without compromising regulatory control, we employed a Triplex RNA motif derived from the mouse non-coding RNA Malat1^35^, combined previously with CRISPR-Cas12a and crRNA for robust DNA-targeting activity^36^. Triplex inclusion to restore closer to optimal Cas13d levels while keeping crRNA expression moderated led to MONARCH 2.0 **(Figure 5A)**, which significantly enhanced the tunability of on-target activity without reintroducing collateral activity **(Figure 5H; Figure S5D)**.

An additional surprising benefit in Vero E6 green monkey cells was successful LP-based gene circuit integration, originally developed for human cells. The AAVS1 orthologous site in the Vero E6 genome had functional sgRNA targeting sites and intact homologous regions, enabling precise Cas9-based genomic LP integration **(Figure S7A)**. The Vero E6 cells engineered with the MONARCH system showed robust capability for targeting viral genomes **(Figure 7D)**, setting the stage for future quantitative human virology studies. Besides MONARCH, we believe that the precisely engineered LP Vero E6 cells can provide a standard, feasible and robust approach to rapidly generate relevant cell line models, to enable genetic screens and other investigations.

In conclusion, the optimized MONARCH circuit we developed precisely controls the expression of Cas13d and crRNA, enabling effective RNA downregulation with minimal collateral activity. By establishing this system in engineered Vero E6 cells, we have illustrated its potential to significantly impact biological research and therapeutic development, particularly in the context of human virology.

## METHODS

### Cell culture

HEK293 cells were originally obtained from American Type Culture Collection (ATCC®). All engineered versions of HEK293 cells were cultured in DMEM media with 10% fetal bovine serum (FBS) and 1% penicillin-streptomycin. The Vero E6/TMPRSS2 cell line was a gift from Dr. Charles M. Rice at Rockefeller University. All engineered versions of Vero E6 cells were cultured in DMEM media with 10% fetal bovine serum (FBS) and 1% penicillin-streptomycin. Both cell lines were maintained regularly at a 2-4 days interval for each passage based on usage demands. All cell lines and their variants were maintained in Panasonic MCO-170AICUVL-PA cellIQ Series CO2 Incubators at 37⁰C and 5% CO2. The cells were used in experiments within 15 passages after their arrival in the laboratory.

### Plasmid Construction

For the generation of the LPutopia-bearing Vero E6 cell line (Vero E6 LP cell), the **LPutopia-8** genome-targeting vector was constructed based on the earlier version LPutopia-7 (Addgene #199212) assembled in the previous studies^12^. To circumvent antibiotic selection conflicts, we replaced the neomycin resistance gene (NeoR), which was previously introduced along with the TMPRSS2 gene into the Vero E6 model cell line, with a puromycin resistance gene (PuroR) in the LP cassette. Briefly, new selection marker puromycin resistance gene (PuroR) was incorporated into eGFP expression cascade using T2A separation, while original neomycin resistance gene (NeoR) was deleted from the cassette. Every other element was remained the same as LPutopia-7.

For constructing the RMCE vectors, we first built **pUt-mNF-Cas13d** circuit plasmid by replacing the GFP reporter with mCherry-P2A-RfxCas13d target cascade, obtained from pSLQ5428_pHR_EF1a-mCherry-P2A-Rfx_Cas13d-2xNLS-3xFLAG (Addgene #155305), in the previous cloned pUt-mNF-GFP RMCE plasmid^12^ (Addgene #199220). Then **MONARCH 1.0** circuit was established by inserting pre-crRNA motif (with GFP-targeting crRNA) right after coding sequences in the 3’UTR region of pUt-mNF-Cas13d plasmid. Further **MONARCH 2.0** circuit was cloned by adding Triplex motif, cloned from pCE048-SiT-Cas12a (Addgene #128124), between coding sequences and pre-crRNA motif. All the all-in-one testing constructs were similarly cloned by inserting SV40-GFP and/or SV40-BFP and/or hU6-2O-crRNA expression cascades into the FRT3/FRT flanking region, to be integrated together with gene circuits via FLP-RMCE simultaneously.

For crRNA plasmids generated in this study, we took the template from pSLQ5429_pUC_hU6-crScaffold_EF1α-BFPv3 (Addgene #155306) and cloned in individual crRNAs using restriction-ligation method. A SV40 replacing EF1α version of BFP-targeting and GFP-targeting crRNA were also cloned. TetR-regulatable hU6 promoter, hU6-2O, was DNA synthesized by IDT with reference sequences taken from previous study^33^, then cloned into BFP- and GFP-targeting crRNA plasmids to replace the original hU6 promoter. While BACH1-targeting crRNA array was combined together while individual hU6 promoter was preserved, SARS-CoV-2-targeting crRNA array was assembled into one expression cascade using golden gate assembly. **MONARCH** system was cloned by assembling pre-crRNA or pre-crRNA array from crRNA plasmids with **pUt-mNF-Cas13d** using NEBuilder® HiFi DNA Assembly.

For constructing SARS-CoV-2 genomic fragment targeting reporter plasmid, we first obtained the CoV-2 genomic fragment from pSLQ5080_pHR_PGK_sfGFP_CoV-F2 (Addgene #155304) and then merged it into the 3’UTR of eGFP expression cascade in LPutopia-7 (Addgene #199212) plasmid, with RFP deleted from the backbone.

The AAVS1-targeting eSpCas9 vector (Addgene #199213) was constructed in the previous study^12^. the codon-optimized FLP recombinase expressing vector-**pCAG-Flpo** also available as Addgene plasmid #60662 was a gift from Dr. Massimo Scanziani. The crRNA template plasmid, SARS-CoV-2-targeting crRNA and non-targeting plasmids, RfxCas13d plasmid and SARS-CoV-2 genomic fragment plasmid were gifts from Dr. Stanley Qi. The high-fidelity RfxCas13d expressing plasmid was available as Addgene plasmid #190034. We used NEBuilder® HiFi DNA Assembly for most of the plasmid cloning. sgRNA and all crRNA sequences used in this study are available in **Table S2**.

### Transfection and Nucleofection

Lipofection was applied for all HEK293-derived cells. Prior to transfection, cells were plated in 6-well plates and grown to ∼80% confluence. Then pCAG-Flpo helper vector and NF-circuit or MONARCH donor vectors were co-transfected at a 1-to-1 ratio with a final mass of 2.5 µg per well. The vectors were first incubated with 3.75 µl of Lipofectamine 3000 (Invitrogen, L3000-015) in OPTI-MEM media (Gibco, 31985062) for 15-30 minutes. The resulting DNA-lipid complex was then pipetted onto the cell culture. Cells were then incubated for at least 24 hours before refreshing media. Appropriate drugs for selection were added 72 hours post-transfection. Drug selection lasted for at least 14 days before FACS cell sorting. We used 10 µg/ml Blasticidin with 10 µg/ml ganciclovir for all the HEK293 engineering cell selection.

For all Vero E6 and its derived cells, Nucleofection was performed based on the manufacturer’s instructions and recommendations. Briefly, newly thawed Vero E6 cells were plated in a T-25 flask and sub-cultured 3-5 days before Nucleofection. Next, cells were harvested by adding trypsin and counted using a Cellometer Auto T4 (Nexcelom Bioscience). Around 1x10^6^ cells were collected and centrifuged at 200 x g for 10 minutes at room temperature. Then the supernatant was removed and cells were resuspended in 100 µl room-temperature Nucleofector^®^ Kit V (Lonza, VCA-1003) solution. The LPutopia-8/espCas9 or pCAG-Flpo/MONARCH donor vector combinations were co-transfected at a 1-to-1 ratio with a final mass of 2 µg per sample. The cell/DNA suspension was transferred into the certified Nucleofector^®^ cuvette and the V-001 program of Nucleofector^®^ 2b Device (Lonza, AAB-1001) was applied. Finally, transfected cells were buffered with fresh media and gently transferred into a freshly prepared 6-well plate. Drug Selection started 24-48 hours post-nucleofection and lasted for at least 14 days before FACS cell sorting. We used 4 µg/ml of puromycin for Vero E6 LP cell selection and 2 µg/ml Blasticidin with 5 µg/ml ganciclovir for Vero E6 MONARCH cell selection.

For all the transient transfection assays, engineered HEK293 cells were seeded and induced 72 hours prior to plasmid transfection using Lipofectamine 3000 as described above, while engineered Vero E6 cells were collected after 72-hour pre-induction for nucleofection with plasmids then plated back into 6-well plates for incubation. For both cases, corresponding Dox induction levels were maintained during plasmid incubation, and 72 hours incubation was typically applied prior to flow cytometry or cell viability assay.

### Flow cytometry

For each sample, cells were cultured for one passage prior to experimentation. Subsequently, approximately 50,000 to 80,000 cells, harvested from T-25 flasks at 80% confluence, were seeded into 24-well plates, maintaining three technical replicates for each concentration of the inducer. Doxycycline (Dox) was added to each well to achieve concentrations ranging from 0.01 to 100 ng/ml. The cells were then incubated for 72 hours before being transferred to a 96-well plate, with a final volume of 250 µl per well. Flow cytometry analysis was conducted using a BD LSRFortessa™ flow cytometer equipped with a High Throughput Sampler (HTS) at the Stony Brook Genomics Core Facility. We collected multiple fluorescence signal data for each Dox concentration, recording at least 10,000 events for stably integrated targets and 50,000 for transiently provided targets within a predefined FSA/SSA gate. GFP intensity was measured using the FITC channel, mCherry intensity via the Texas-Red channel, and BFP intensity from the Pacific-Blue channel, maintaining identical PMT voltage settings across all samples and induction levels.

### RNA isolation and RT-qPCR

For BACH1 downregulation RT-qPCR test, 100-300k HEK cells were first pre-induced for 72 hours with each Dox concentration in 6-well plates prior to BACH1-targeting crRNA transfection. 144 hours (6-day) after transfection, RNA was then isolated using the RNeasy Plus Mini Kit (Qiagen, 74134). 1 µg of total RNA from each sample was converted to cDNA using iScript™ Reverse Transcription Supermix (Bio-Rad, 1708841). Then qPCR reactions were set up using TaqMan^®^ Fast Advanced Master Mix (ThermoFisher Scientific, 4444557) with TaqMan^®^ Gene Expression Assay and run using the QuantStudio™ 3 Real-Time PCR System (Eppendorf, A28137) at standard curve mode, using the BACH1 TaqMan^®^ probe (ThermoFisher Scientific, Hs00230917_m1) and GAPDH reference probe (ThermoFisher Scientific, 4326317) in a multiplexing setup. Eventually, BACH1 mRNA level quantitation was calculated with reference to the GAPDH level in three independent repeats.

### Cell viability assay

For cell viability assays, cells were Dox-induced 72 hours before seeding and 3,000 – 5,000 cells of each induction condition were plated in 96-well plates with 4 replicates with Dox level maintained. Around 6 hours after seeding, transfection of corresponding plasmid or plasmid combinations were applied into settled cells in 96-well plates. all the seeded wells were assayed using alamarBlue™ HS Cell Viability Reagent (Invitrogen, A50100) for viable cells 72 hours post-transfection with dox level maintained throughout. To measure multi-day proliferation, total 6 replicates were seeded at the beginning and then 3 replicates were assessed every 24 hours.

Cells were incubated in alamarBlue reagent for 4 hours and then taken for absorbance measurement at wavelengths of 570 nm and 600 nm with media blank control using Tecan Infinite Pro 200 (Tecan U.S. INC) spectrophotometer. Each absorbance value was adjusted by subtracting the media blank absorbance at the same wavelength. Cell proliferation was measured as the alamarBlue reduction score (S) calculated as:

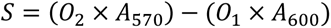

Where 𝑂_1_ and 𝑂_2_ are the molar extinction coefficients of oxidized alamarBlue at 570 nm and 600 nm, respectively; and 𝐴_570_ and 𝐴_600_ are the absorbances of test wells at 570 nm and 600 nm, respectively. Relative cell viability was then calculated as the relative differences between the average scores of the treated wells to untreated control wells.

### Data processing and statistical analysis

Flow cytometry data were analyzed with the FlowJo^®^ software version 10 (Becton, Dickinson and Company). Forward-scatter and side-scatter gates were predefined for each cell type or assay based on the reference sample pretests to exclude debris. Also, a fluorescence-based gate was imposed for FACS cell sorting for desired target cells. Most of the data plots as well as statistical analysis were generated and performed using GraphPad Prism 8 (GraphPad Software). Statistical details are in the figure legends, including the statistical tests used. In all figures, results were presented as means ± S.D. unless otherwise noted in the figure legend. *P < 0.05 was considered statistically significant, indicated by “**\***” in the figure legend.

### Computational Modeling and Mathematical Derivations

For each system (i.e. mNF-Cas13d and MONARCH), we wrote the corresponding series of biochemical reactions. From these reactions, systems of ordinary differential equations were established. These equations were set equal to zero to solve for the steady state values of each chemical species. To simulate inducible Cas13d expression, we adjusted the rate of Cas13d mRNA transcription (sR) over several orders of magnitude. Algebraic equations were solved with the help of Mathematica (version 14.0.0.0), and solutions were plotted and visualized using MATLAB (version R2022a). See the **Supplemental Results** for detailed information about the model equations and parameters used.

## Supporting information

Supplementary Results, Figures and Tables

## ACKNOWLEDGMENTS

We thank Dr. Feng Zhang for sharing espCas9, Dr. Massimo Scanziani for sharing the Flp recombinase and Dr. Charles M. Rice for Vero E6 cell line. We thank S. Lei Qi for the SARS-CoV-2 fragments, RfxCas13d and the corresponding crRNAs. We thank the Balázsi lab members and Dr. Erich Mackow for insightful discussions and comments. GB was supported by the National Institutes of Health, NIGMS MIRA Program (R35 GM122561) and by the Laufer Center for Physical and Quantitative Biology. GB is also grateful for support by a Stony Brook University Office of the Vice President for Research Seed Grant and a Stony Brook Cancer Center Engineering, Physical Sciences and Oncology Pilot Funds.

## AUTHOR CONTRIBUTIONS

Conceptualization, Y.W., and G.B.; Formal Analysis, Y.W. and G.B.; Methodology, Y.W., and G.B.; Investigation, Y.W., D.C., C.H., and G.B.; Data Curation and Visualization, Y.W. C.H. and G.B.; Mathematical Modeling, C.H. and G.B.; Writing –Original Draft, Y.W. and G.B.; Writing – Review & Editing, Y.W., C.H. and G.B.; Funding Acquisition and Resources, G.B.; Supervision, G.B.; Project Administration, G.B.

## DATA AND CODE AVAILABILITY

All data is available at: Balázsi Lab Wiki on OpenwetWare, Data+Plasmids+Software. https://openwetware.org/wiki/CHIP:Data. Deposited 22 August 2024.

All materials, code and data will also be available by the lead contact upon request. Additionally, we deposit some of the plasmids generated from this study with Addgene.

