## Supplementary Results, Figures and Tables for "Optimizing a CRISPR-Cas13d gene circuit for tunable target RNA downregulation with minimal collateral RNA cutting"

**Model Description and Reactions**

The species modeled are shown in **Table S3**.

| Species | Description |
|---|---|
| G | Guide RNA |
| R | Cas13d mRNA transcript |
| C | Cas13d protein |
| T | Target RNA |
| N | Nonspecific RNA |
| A | Activated Cas13d complex |
| H | Hyperactive Cas13d complex |
| U | Unstable Cas13d mRNA transcript |

**Table S3**: Description of species modeled.

*<u>mNF-Cas13d Set-up</u>*

First, we consider the mNF-Cas13d system, which can be composed of the following reactions:

$$\emptyset \xrightarrow{s_G} G \xrightarrow{g_G} \emptyset$$

$$\emptyset \xrightarrow{s_R} R \xrightarrow{g_R} \emptyset$$

$$R \xrightarrow{s_C} R + C$$

$$\emptyset \xrightarrow{s_T} T \xrightarrow{g_T} \emptyset$$

$$\emptyset \xrightarrow{s_N} N \xrightarrow{g_N} \emptyset$$

$$C + G \xrightarrow{a} A$$

$$A + T \xrightarrow{h} H$$

$$H + T \xrightarrow{c_T} H$$

$$H + N \xrightarrow{c_N} H$$

$$C \xrightarrow{g_C} \emptyset$$

$$A \xrightarrow{g_A} \emptyset$$

$$H \xrightarrow{g_H} \emptyset$$

In summary, the guide RNA (G) is produced from a separate promoter. This can then associate with the Cas13d protein to form an active complex (A). The active complex can then start cutting target RNA (T) to become a “hyperactive complex” (H). This hyperactive complex can then cut both target RNA (T) and nonspecific RNA (N). See Table S4 for descriptions of the parameters used.

From these reactions, the corresponding set of ordinary differential equations (ODEs) can be written as the following (and set equal to zero to find the corresponding steady states):

$$\dot{G} = s_G - g_G G - aCG = 0$$

$$\dot{R} = s_R - g_R R = 0$$

$$\dot{C} = s_C R - g_C C - aCG = 0$$

$$\dot{T} = s_T - g_T T - hAT - c_T HT = 0$$

$$\dot{N} = s_N - g_N N - c_N HN = 0$$

$$\dot{A} = aCG - hAT - g_A A = 0$$

$$\dot{H} = hAT - g_H H = 0$$

First, we consider the first three equations:

$$R = s_R / g_R$$

$$G = s_G / (g_G + aC)$$

$$s_C \left(\frac{s_R}{g_R}\right) - g_C C - aC\left(\frac{s_G}{g_G + aC}\right) = 0$$

This can be rearranged into the following quadratic:

$$[ag_C]C^2 + \left[as_G + g_C g_G - as_C \frac{s_R}{g_R}\right] C - g_G s_C \frac{s_R}{g_R} = 0$$

Which can be solved using the positive term of the quadratic formula to solve for C in terms of the necessary parameters. Namely,

$$C = \frac{-\left(as_G + g_C g_G - as_C \frac{s_R}{g_R}\right) + \sqrt{(as_G + g_C g_G - as_C \frac{s_R}{g_R})^2 - 4ag_C g_G s_C \frac{s_R}{g_R}}}{2ag_C}$$

Note that once C is solved, this can also be used to solve for G. Also note that we can substitute $\eta = aCG$, since C and G are solved from the previous equations, so $\eta$ is a positive constant.

Next, we consider the target RNA dynamics equations (namely, $\dot{T}, \dot{A}, \dot{H}$):

$$H = hAT / g_H$$

$$A = \eta / (hT + g_A)$$

$$s_T - g_T T - hAT - c_T HT = 0$$

Note that $c_T \propto \frac{H}{K+H}$ (see Table S4), which makes the equations more complicated. In general, the solution will be the positive root of the corresponding cubic equation. Mathematica (version 14.0.0.0) was used to solve this equation for T with the corresponding parameters.

Similarly, once T is solved, it can be used to solve for the other species in the system using the equations above. Additionally, N can also be solved with the following equation:

$$N = s_N/(c_N H + g_N)$$

*MONARCH Set-Up*

Next, the MONARCH system can be described with the following reactions:

$$\emptyset \xrightarrow{s_R} R \xrightarrow{g_R} \emptyset$$

$$R \xrightarrow{s_C} R + C$$

$$U \xrightarrow{s_C} U + C$$

$$\emptyset \xrightarrow{s_T} T \xrightarrow{g_T} \emptyset$$

$$\emptyset \xrightarrow{s_N} N \xrightarrow{g_N} \emptyset$$

$$C + R \xrightarrow{a} A + U$$

$$A + T \xrightarrow{h} H$$

$$H + T \xrightarrow{c_T} H$$

$$H + N \xrightarrow{c_N} H$$

$$C \xrightarrow{g_C} \emptyset$$

$$A \xrightarrow{g_A} \emptyset$$

$$H \xrightarrow{g_H} \emptyset$$

$$U \xrightarrow{g_U} \emptyset$$

In summary, the guide RNA is part of the Cas13d mRNA transcript (R). Thus, Cas13d processes its own guide, creating an unstable mRNA transcript (U), which can still express Cas13d, but has increased degradation due to lacking a polyA tail. Similarly, this active complex can then start cutting target RNA (T) to become a "hyperactive complex" (H). This hyperactive complex can then cut both target RNA (T) and nonspecific RNA (N). See Table S4 for descriptions of the parameters used.

From these reactions, the corresponding set of ordinary differential equations (ODEs) can be written as the following (and set equal to zero to find the corresponding steady states):

$$\dot{U} = aCR - g_U U = 0$$

$$\dot{R} = s_R - g_R R - aCR = 0$$

$$\dot{C} = s_C(U + R) - g_C C - aCR = 0$$

$$\dot{T} = s_T - g_T T - hAT - c_T HT = 0$$

$$\dot{N} = s_N - g_N N - c_N HN = 0$$

$$\dot{A} = aCR - hAT - g_A A = 0$$

$$\dot{H} = hAT - g_H H = 0$$

First, we consider the first three equations:

$$U = \frac{aCR}{g_U} = (s_R - g_R R)/g_U$$

$$C = (s_R - g_R R)/aR$$

$$s_C((s_R - g_R R)/g_U + R) - g_C((s_R - g_R R)/aR) - a((s_R - g_R R)/aR)R = 0$$

This can be rearranged into the following quadratic:

$$\left[as_C\left(1 - \frac{g_R}{g_U}\right) + ag_R\right]R^2 + \left[\frac{as_C s_R}{g_U} + g_C g_U - as_R\right]R - g_C s_R = 0$$

Which can be solved using the positive term of the quadratic formula to solve for R in terms of the necessary parameters. Namely,

$$R = \frac{-\left(\frac{as_C s_R}{g_U} + g_C g_U - as_R\right) + \sqrt{(\frac{as_C s_R}{g_U} + g_C g_U - as_R)^2 + 4(as_C\left(1 - \frac{g_R}{g_U}\right) + ag_R)(g_C s_R)}}{2(as_C\left(1 - \frac{g_R}{g_U}\right) + ag_R)}$$

Note that once R is solved, this can also be used to solve for G and C. Also note that we can substitute $\eta = aCR$, since C and R are solved from the previous equations, so $\eta$ is a positive constant.

Now, with the substitution of $\eta$, the equations for T, A, H, and N take the same form as the mNF-Cas13d model. Therefore, from the structure of the steady state equations, the difference is in the $\eta$ parameter (in other words, the synthesis term from the active complex).

*Parameters*

In order to capture the differences between these two systems that we observed experimentally **(Fig S6A)**, we also adjusted key parameters accordingly **(Table S4)**.

First, to capture the nonlinear dynamics of target RNA and nonspecific RNA cutting, we made the cutting terms nonlinear. In particular, we made the cutting rate of target RNA ($c_T$) follow Michaelis Menten kinetics, because of the spatial activation of the *trans* cleavage of Cas13d by target cutting needing to occur first before nonspecific RNA cutting can be observed. Additionally, we made the cutting rate of nonspecific RNA ($c_N$) follow hill equations, to capture the phenomenon of the delay in cutting of nonspecific RNA (again, due to the delay in activation based on co-localization needed for target RNA cutting).

To further emphasize the difference in these two models, we adjusted the rates of cutting target RNA and nonspecific RNA ($h, c_T, c_N$). In particular, we set the target RNA cutting rates to be higher in the MONARCH model compared to the mNF model, based on the limited amount of activated Cas13d complexes in comparison to the total target RNA pool (see **Fig 6A**). Accordingly, we set the nonspecific RNA cutting rates to be higher in the mNF model for the same reason.

| Parameter | Description | Value in mNF Model | Value in MONARCH Model |
|---|---|---|---|
| $s_G$ | Synthesis rate of guide RNA (G) | 100 | N/A |
| $s_R$ | Synthesis rate of Cas13d mRNA (R) | Ranges from 0.001 to 1000 | Ranges from 0.001 to 1000 |
| $s_T$ | Synthesis rate of target RNA (T) | 1 | 1 |
| $s_N$ | Synthesis rate of nonspecific RNA (N) | 1 | 1 |
| $s_C$ | Translation rate of Cas13d protein (C) | 1 | 1 |
| $g_G$ | Degradation rate of guide RNA (G) | 0.1 | 0.1 |
| $g_R$ | Degradation rate of Cas13d mRNA (R) | 0.1 | 0.1 |
| $g_T$ | Degradation rate of target RNA (T) | 0.1 | 0.1 |
| $g_N$ | Degradation rate of nonspecific RNA (N) | 0.1 | 0.1 |
| $g_U$ | Degradation rate of unstable Cas13d mRNA (U) | N/A | 1 |
| $g_C$ | Degradation rate of Cas13d protein (C) | 0.01 | 0.01 |
| $g_A$ | Degradation rate of active Cas13d complex (A) | 0.01 | 0.01 |
| $g_H$ | Degradation rate of hyperactive Cas13d complex (H) | 0.01 | 0.01 |

| | | | |
|---|---|---|---|
| a | Active Cas13d complex (A) formation rate | 1 | 1 |
| h | Cutting rate of target RNA (T) by active Cas13d complex (A) | 1 | 10 |
| $c_T$ | Cutting rate of target RNA (T) by hyperactive Cas13d complex (H) | $1 * \frac{H}{10 + H}$ | $10 * \frac{H}{10 + H}$ |
| $c_N$ | Cutting rate of nonspecific RNA (N) by hyperactive Cas13d complex (H) | $10 * \frac{H^2}{10^2 + H^2}$ | $1 * \frac{H^2}{100^2 + H^2}$ |

**Table S4:** Description and values of parameters in the models. See text for justification of parameter differences between models. N/A = not applicable (parameter is not in that model).

To simulate each model, we adjusted the synthesis rate of Cas13d mRNA ($s_R$) uniformly (in log space) from 0.001 to 1000. We then solved the algebraic equations using the solutions presented and the parameters listed, and plotted the solutions using MATLAB (version r2022a).

To get the initial amounts of target RNA and nonspecific RNA, we considered the reactions without any Cas13d cutting dynamics. Accordingly, the ODEs simplify to give the results $T = s_T/g_T$ and $N = s_N/g_N$.

To investigate the effect of target RNA cutting by hyperactive Cas13d, we set $c_T = 0$, and kept all other parameters the same. Furthermore, to estimate the target RNA efficiency, we calculated the average slope along the dose response range. In particular, we took the average slope from the basal level (the point where the curve starts rapidly decreasing) to the saturating point (where the curve drastically changes slope for higher levels of $s_R$.

To simulate the effect of hfCas13d, we only considered the mNF model. We decreased the cutting rate of target RNA by 10 (in particular, we set $h = 0.1$, and $c_T = 0.1 * \frac{H}{10+H}$), and we set the rate of nonspecific RNA cutting to zero (i.e. $c_N = 0$).

### Supplementary Figures

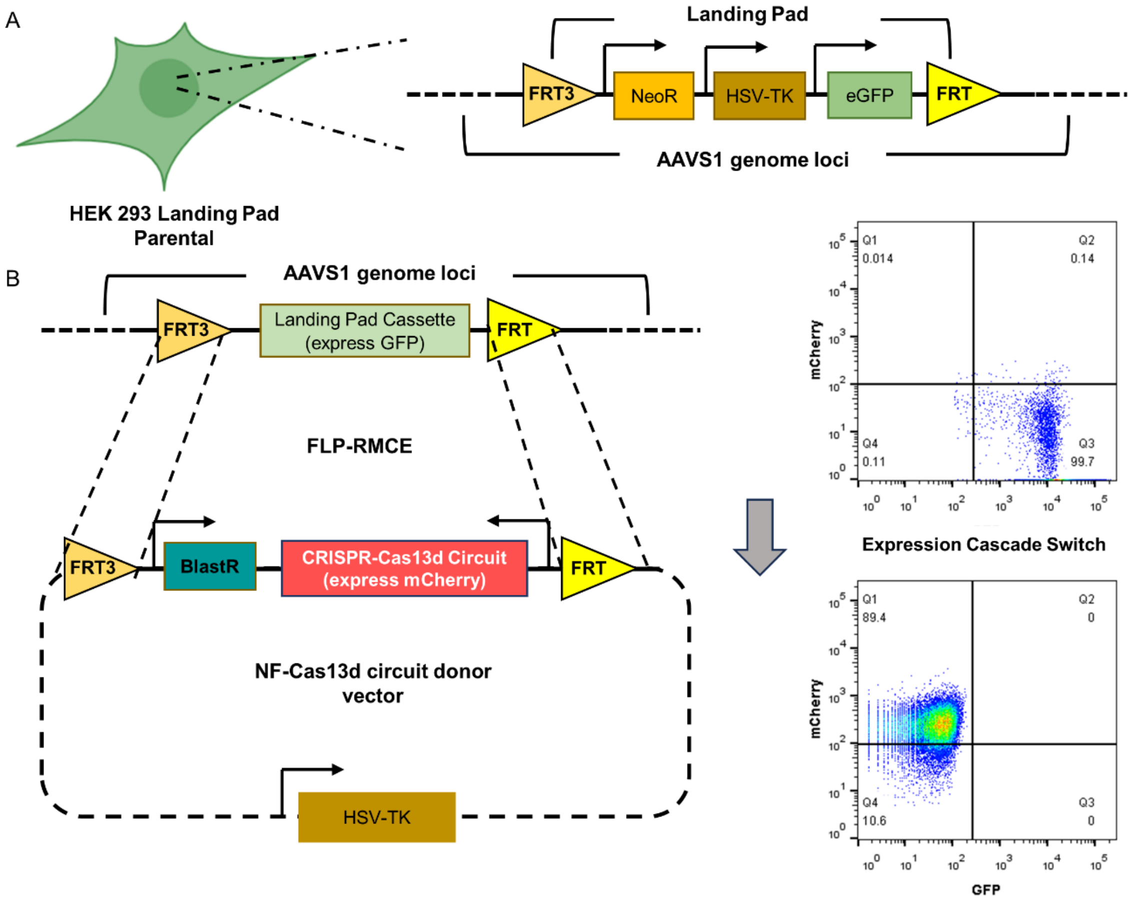

**Figure S1. Integration of negative-feedback CRISPR-Cas13d gene circuit with landing-pad platform and recombinase-mediated cassette exchange.**

**(A)** Diagram of engineered HEK 293 cell with a single-copy landing pad locating in the AAVS1 safe harbor site. FRT, FRT3: Flp-recombinase target sites; NeoR: Neomycin Resistance Gene; HSV-TK: Herpes Simplex Virus (HSV) thymidine kinase (TK); eGFP: enhanced Green Fluorescence Protein.

**(B) Left panel:** Schematic diagram of repeatable AAVS1 site-specific integration of genetic payloads such as CRISPR-Cas13d negative-feedback (NF) circuit through Flp-recombinase-mediated cassette exchange (FLP-RMCE). **Right panel:** RMCE-based integration will result in a DNA construct switching between donor plasmid and target site. Successful integration of mNF-Cas13d gene circuit showed a fluorescence signal switch from green (eGFP) to red (mCherry). HSV-TK in the donor plasmid backbone serves as a negative selection marker against random integration.

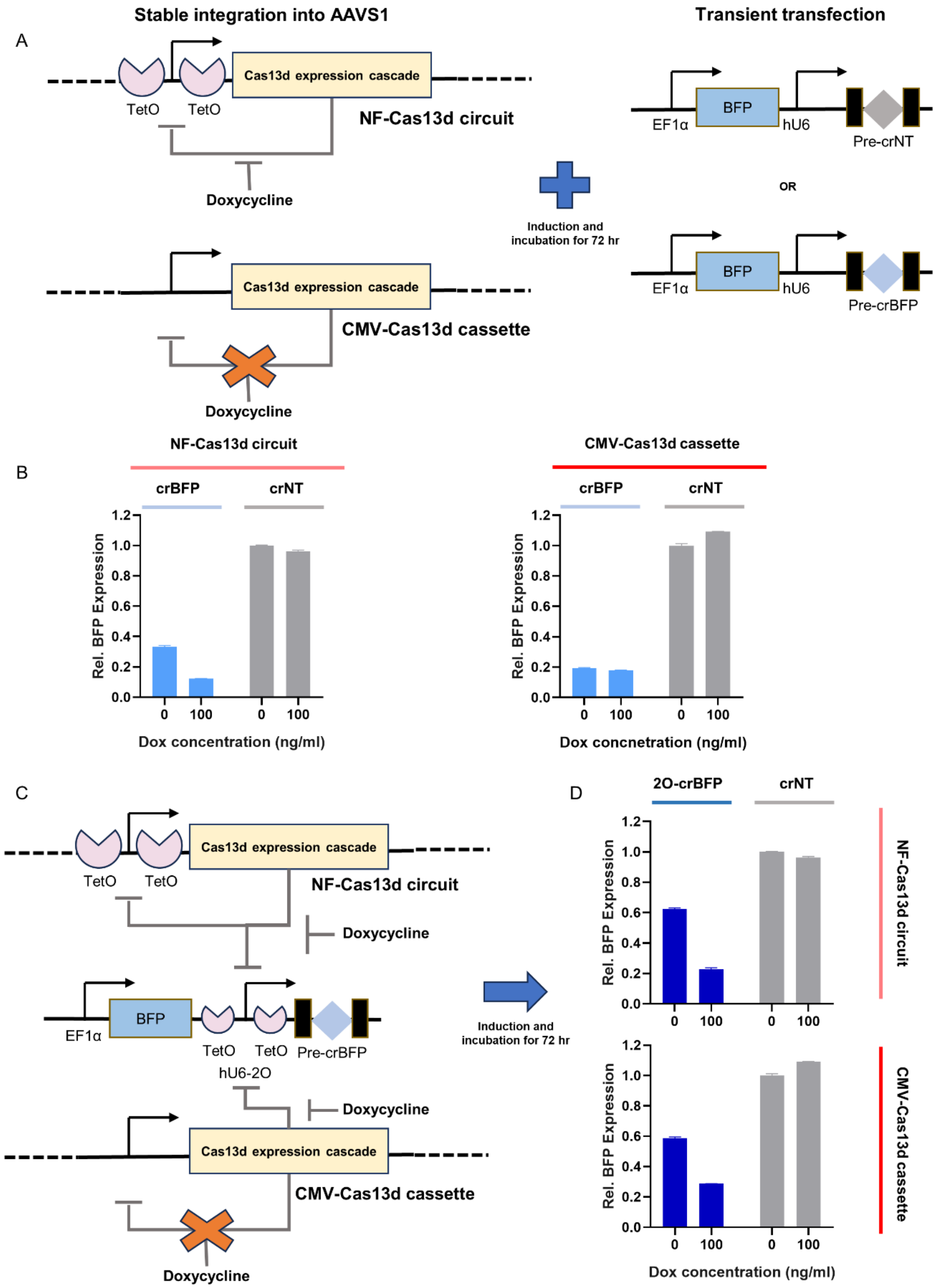

A
Stable integration into AAVS1
Transient transfection
Cas13d expression cascade
TetO
TetO
NF-Cas13d circuit
Doxycycline
CMV-Cas13d cassette
Induction and incubation for 72 hr
BFP
EF1α
hU6
Pre-crNT
OR
Pre-crBFP
B
NF-Cas13d circuit
CMV-Cas13d cassette
crBFP
crNT
Rel. BFP Expression
Dox concentration (ng/ml)
Dox concnetration (ng/ml)
C
hU6-2O
D
2O-crBFP

**Figure S2. Cas13d's basal target reduction depends on crRNA abundance.**

**(A)** Diagram of single-vector transfection to test Cas13d inducible on-target activity. Plasmid encoding BFP reporter and BFP targeting guide or non-targeting guide were transfected into engineered HEK 293 cells with or without doxycycline (Dox) induction and incubated for 72 hours before flow cytometry. Compared to mNF-Cas13d circuit, CMV-Cas13d cassette only removed the two Tetracycline Operator (TetO2) sites from CMV promoter without any changes in the Cas13d expression cascade.

**(B)** Comparison of BFP reduction induced by Cas13d and unregulated crRNA targeting 72 hours post-transfection with or without doxycycline induction in both engineered HEK 293 cells. All the BFP expression levels were normalized to uninduced sample with non-targeting guide, n=3.

**(C)** Diagram of single-vector transfection to test Cas13d inducible on-target activity with regulated crRNA expression. crRNA targeting BFP was expressed from a synthetically modified human U6 (hU6) promoter containing TetO2 sites flanking the TATA-box. hU6-2O promoter can be then repressed by Tetracycline Repressor (TetR) expressed from mNF-Cas13d circuit. CMV-Cas13d cassette can still regulate the hU6-2O promoter due to the intact expression of TetR.

**(D)** Comparison of BFP reduction induced by Cas13d and regulated crRNA targeting 72 hours post-transfection with or without doxycycline induction in both engineered HEK 293 cells. All the BFP expression levels were normalized to uninduced sample with non-targeting guide, n=3.

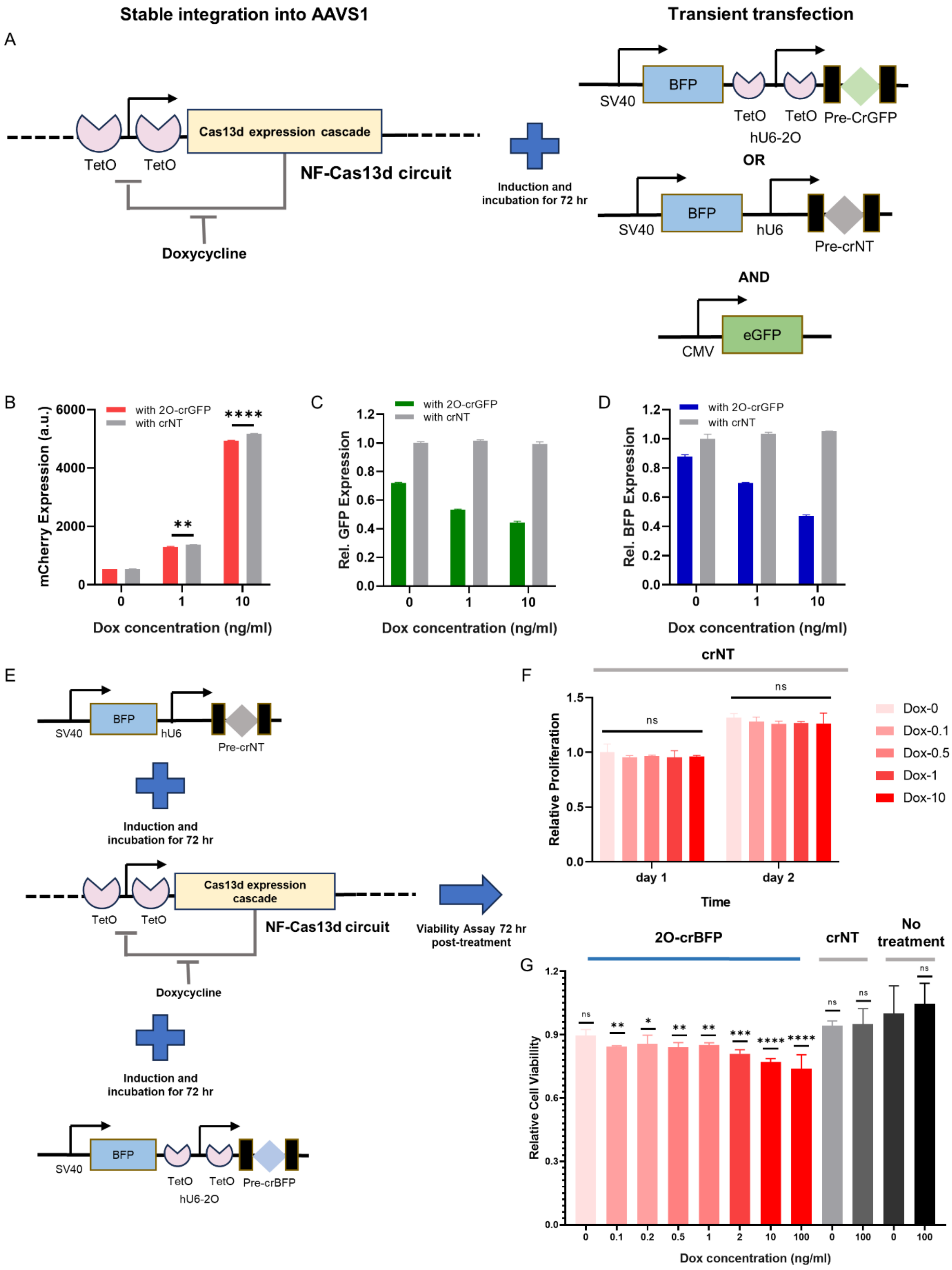
Stable integration into AAVS1
Transient transfection
A
TetO
TetO
Cas13d expression cascade
NF-Cas13d circuit
Doxycycline
Induction and incubation for 72 hr
SV40
BFP
TetO
TetO
hU6-2O
Pre-CrGFP
OR
SV40
BFP
hU6
Pre-crNT
AND
CMV
eGFP
B
with 2O-crGFP
with crNT
mCherry Expression (a.u.)
Dox concentration (ng/ml)
C
Rel. GFP Expression
Dox concentration (ng/ml)
D
Rel. BFP Expression
Dox concentration (ng/ml)
E
SV40
BFP
hU6
Pre-crNT
Induction and incubation for 72 hr
Cas13d expression cascade
NF-Cas13d circuit
Doxycycline
Viability Assay 72 hr post-treatment
SV40
BFP
TetO
TetO
hU6-2O
Pre-crBFP
F
crNT
Relative Proliferation
day 1
day 2
Time
Dox-0
Dox-0.1
Dox-0.5
Dox-1
Dox-10
G
2O-crBFP
crNT
No treatment
Relative Cell Viability
Dox concentration (ng/ml)

**Figure S3. Cas13d on-target activity induces dose-dependent collateral damage.**

**(A)** Diagram of experimental setup to assess collateral damage from activated Cas13d with regulated crRNA expression. Target GFP expressed from CMV promoter was co-transfected with either TetR-regulated GFP-targeting crRNA, or non-targeting crRNA. Both crRNA plasmids contain the same BFP gene expressed from SV40 promoter. Cells were transfected while induced 72 hours before flow cytometry.

**(B)** Comparison of mCherry dose-responses from mNF-Cas13d circuit 72 hours post-transfection with both co-transfection setups. unpaired two-tailed t-test, n=3, **P<0.01, ****P<0.0001.

**(C)** Comparison of relative GFP levels indicating on-target activity dose-responses from mNF-Cas13d circuit 72 hours post-transfection with both co-transfection setups. All the GFP expression levels were normalized to uninduced sample with non-targeting guide, n=3.

**(D)** Comparison of relative BFP levels indicating off-target activity dose-responses from mNF-Cas13d circuit 72 hours post-transfection with both co-transfection setups. All the BFP expression levels were normalized to uninduced sample with non-targeting guide, n=3.

**(E)** Diagram of experimental setup to test Cas13d induced collateral damage on cell viability. Cells were transfected with single-vector targeting construct to activate Cas13d RNase activity. Cell viability was assessed 72 hours post-treatment.

**(F)** Relative cell viability with non-targeting crRNA transfected into HEK 293 cells with stably integrated mNF-Cas13d circuit. All cell viability measurements were normalized to uninduced samples of each day. One-way ANOVA tests, n=3.

**(G)** Relative cell viability with targeting and non-targeting crRNA transfected into HEK 293 cells with stably integrated mNF-Cas13d circuit. All cell viability measurements were normalized to uninduced no-treatment sample. One-way ANOVA with Tukey's multiple comparisons tests between each dose and uninduced no-treatment control, n=3.

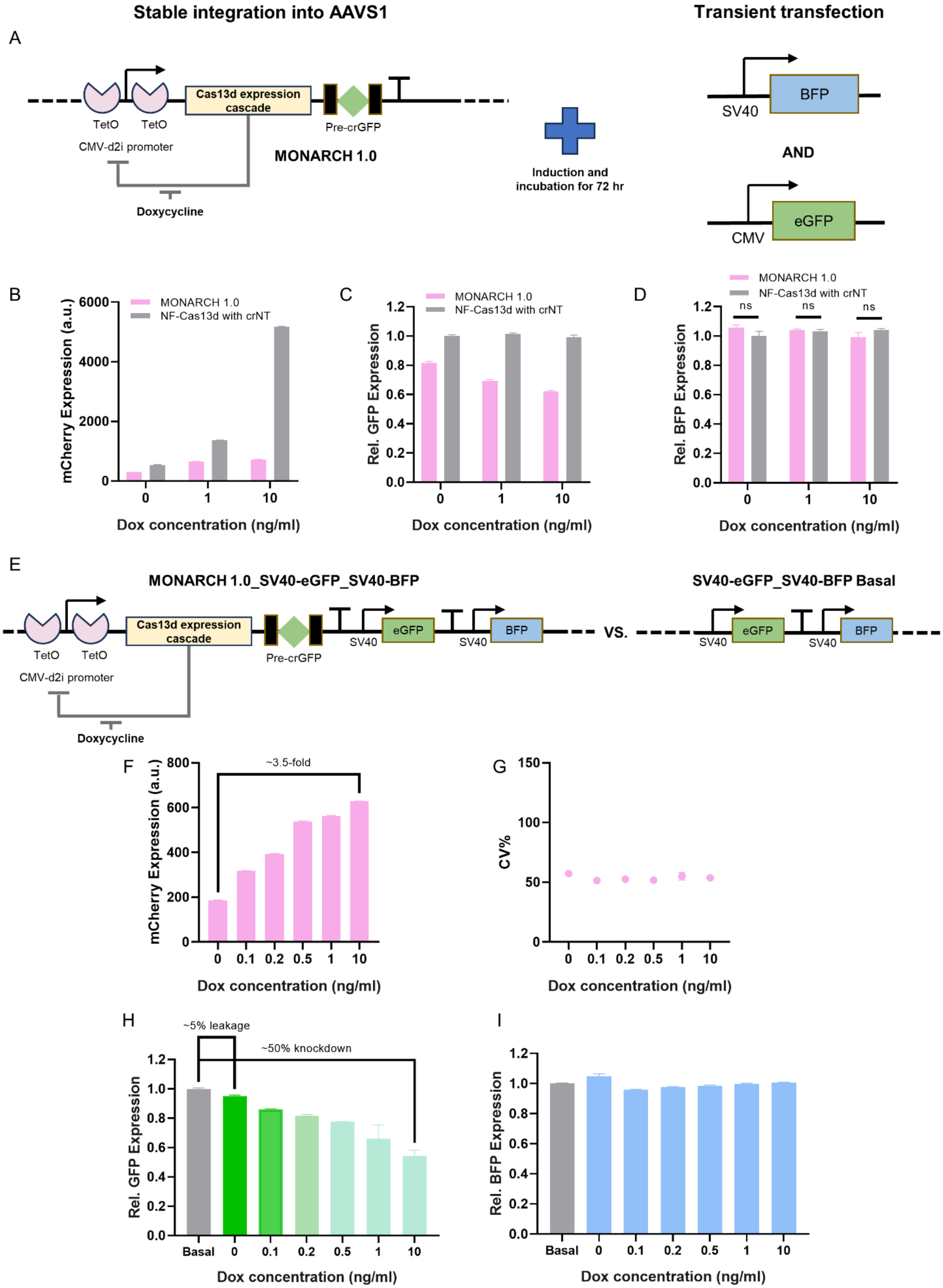
Stable integration into AAVS1
Transient transfection
A
TetO
TetO
CMV-d2i promoter
Cas13d expression cascade
Pre-crGFP
MONARCH 1.0
Doxycycline
Induction and incubation for 72 hr
SV40
BFP
AND
CMV
eGFP
B
mCherry Expression (a.u.)
MONARCH 1.0
NF-Cas13d with crNT
Dox concentration (ng/ml)
C
Rel. GFP Expression
D
Rel. BFP Expression
ns
E
MONARCH 1.0_SV40-eGFP_SV40-BFP
SV40-eGFP_SV40-BFP Basal
VS.
F
~3.5-fold
G
CV%
H
~5% leakage
~50% knockdown
Basal
I

**Figure S4. Multi-level negative autoregulation of Cas13d and crRNA reduces Cas13d collateral damage.**

**(A)** Diagram of experimental setup to assess collateral damage from activated Cas13d with MONARCH 1.0 system. Target GFP expressed from CMV promoter was co-transfected with BFP reporter expressed from SV40 promoter. mNF-Cas13d integrated cells co-transfected with non-targeting crRNA and target GFP was used as control. Cells were transfected while induced 72 hours before flow cytometry.

**(B)** Comparison of mCherry dose-responses between MONARCH 1.0 and mNF-Cas13d circuit 72 hours post-transfection.

**(C)** Comparison of relative GFP levels indicating on-target activity dose-responses from MONARCH 1.0 72 hours post-transfection. All the GFP expression levels were normalized to uninduced mNF-Cas13d sample with non-targeting guide, n=3.

**(D)** Comparison of relative BFP levels indicating off-target activity dose-responses from MONARCH 1.0 72 hours post-transfection. All the BFP expression levels were normalized to uninduced mNF-Cas13d sample with non-targeting guide, n=3.

**(E)** Diagram to assess off-target activity of Cas13d expressed from MONARCH 1.0 on the stably expressing targets from the genome. The whole construct is integrated via FLP-RMCE process using the same HEK 293 Landing pad parentals. Basal level is determined with integration of GFP target and BFP reporter expressed from separate SV40 promoter.

**(F)** Dose-responses of mean fluorescence intensity of mCherry reporter for stably integrated MONARCH 1.0_SV40-GFP_SV40-BFP construct in HEK 293 cells (n=3).

**(G)** Dose-responses of coefficient of variation (CV) of mCherry reporter for stably integrated MONARCH 1.0_SV40-GFP_SV40-BFP in HEK 293 cells (n=3).

**(H)** Dose-responses of relative GFP levels indicating on-target activity for stably integrated MONARCH 1.0_SV40-GFP_SV40-BFP construct in HEK 293 cells. All the GFP expression levels were normalized to the basal sample, n=3.

**(I)** Dose-responses of relative BFP levels indicating off-target activity for stably integrated MONARCH 1.0_SV40-GFP_SV40-BFP construct in HEK 293 cells. All the BFP expression levels were normalized to the basal sample, n=3.

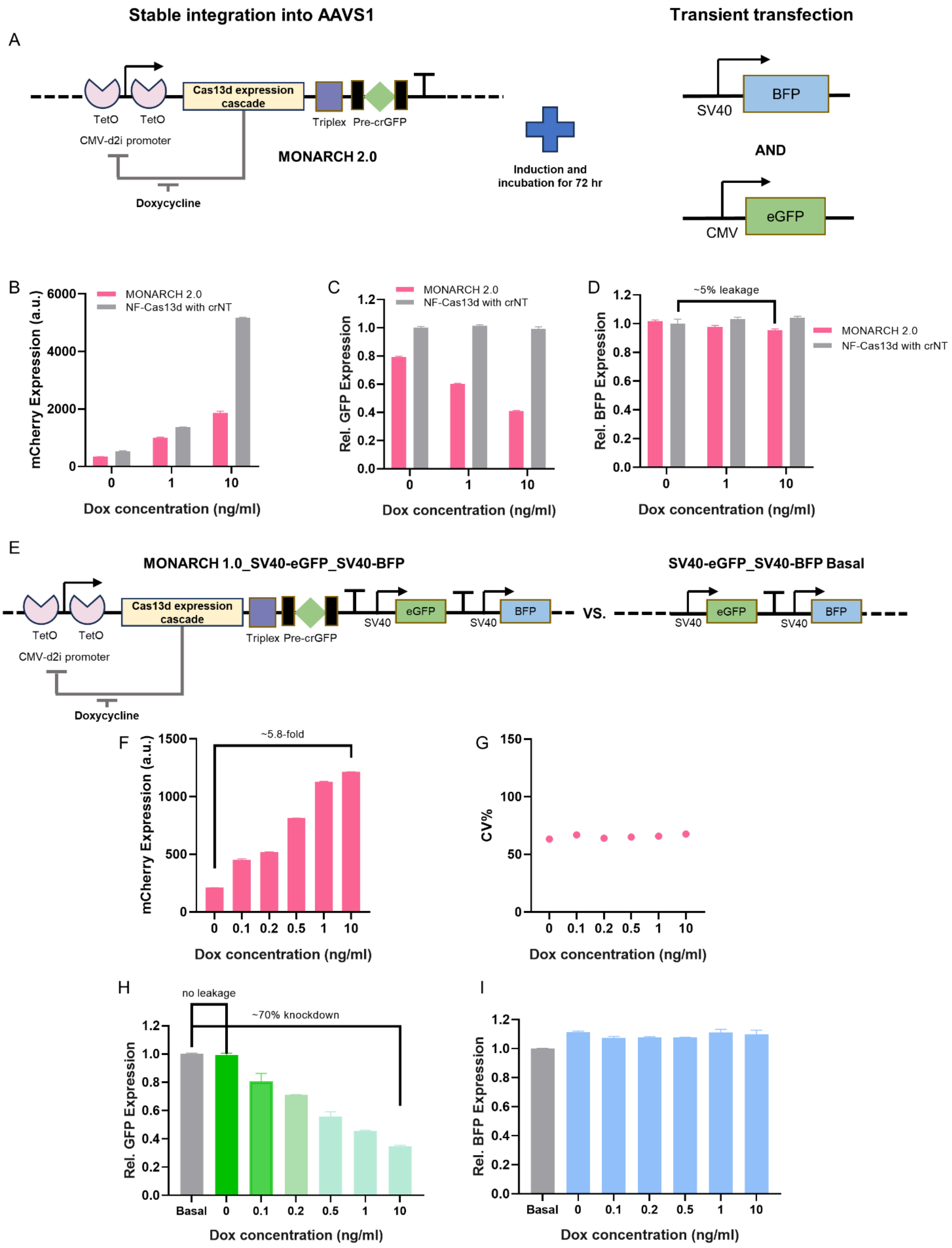
Stable integration into AAVS1
Transient transfection
A
TetO
TetO
CMV-d2i promoter
Cas13d expression cascade
Triplex
Pre-crGFP
Doxycycline
MONARCH 2.0
Induction and incubation for 72 hr
SV40
BFP
AND
CMV
eGFP
B
mCherry Expression (a.u.)
MONARCH 2.0
NF-Cas13d with crNT
Dox concentration (ng/ml)
C
Rel. GFP Expression
D
~5% leakage
Rel. BFP Expression
E
MONARCH 1.0_SV40-eGFP_SV40-BFP
SV40-eGFP_SV40-BFP Basal
VS.
F
~5.8-fold
G
CV%
H
no leakage
~70% knockdown
Basal
I

**Figure S5. Optimized multi-level negative autoregulation of Cas13d and crRNA minimizes Cas13d collateral damage.**

**(A)** Diagram of experimental setup to assess collateral damage from activated Cas13d with MONARCH 2.0 system. Target GFP expressed from CMV promoter was co-transfected with BFP reporter expressed from SV40 promoter. mNF-Cas13d integrated cells co-transfected with non-targeting crRNA and target GFP was used as control. Cells were transfected while induced 72 hours before flow cytometry.

**(B)** Comparison of mCherry dose-responses between MONARCH 2.0 and mNF-Cas13d circuit 72 hours post-transfection.

**(C)** Comparison of relative GFP levels indicating on-target activity dose-responses from MONARCH 2.0 72 hours post-transfection. All the GFP expression levels were normalized to uninduced mNF-Cas13d sample with non-targeting guide, n=3.

**(D)** Comparison of relative BFP levels indicating off-target activity dose-responses from MONARCH 2.0 72 hours post-transfection. All the BFP expression levels were normalized to uninduced mNF-Cas13d sample with non-targeting guide, n=3.

**(E)** Diagram to assess off-target activity of Cas13d expressed from MONARCH 2.0 on the stably expressing targets from the genome. The whole construct is integrated via FLP-RMCE process using the same HEK 293 Landing pad parentals. Basal level is determined with integration of GFP target and BFP reporter expressed from separate SV40 promoter.

**(F)** Dose-responses of mean fluorescence intensity of mCherry reporter for stably integrated MONARCH 2.0_SV40-GFP_SV40-BFP construct in HEK 293 cells (n=3).

**(G)** Dose-responses of coefficient of variation (CV) of mCherry reporter for stably integrated MONARCH 2.0_SV40-GFP_SV40-BFP in HEK 293 cells (n=3).

**(H)** Dose-responses of relative GFP levels indicating on-target activity for stably integrated MONARCH 2.0_SV40-GFP_SV40-BFP construct in HEK 293 cells. All the GFP expression levels were normalized to the basal sample, n=3.

**(I)** Dose-responses of relative BFP levels indicating off-target activity for stably integrated MONARCH 2.0_SV40-GFP_SV40-BFP construct in HEK 293 cells. All the BFP expression levels were normalized to the basal sample, n=3.

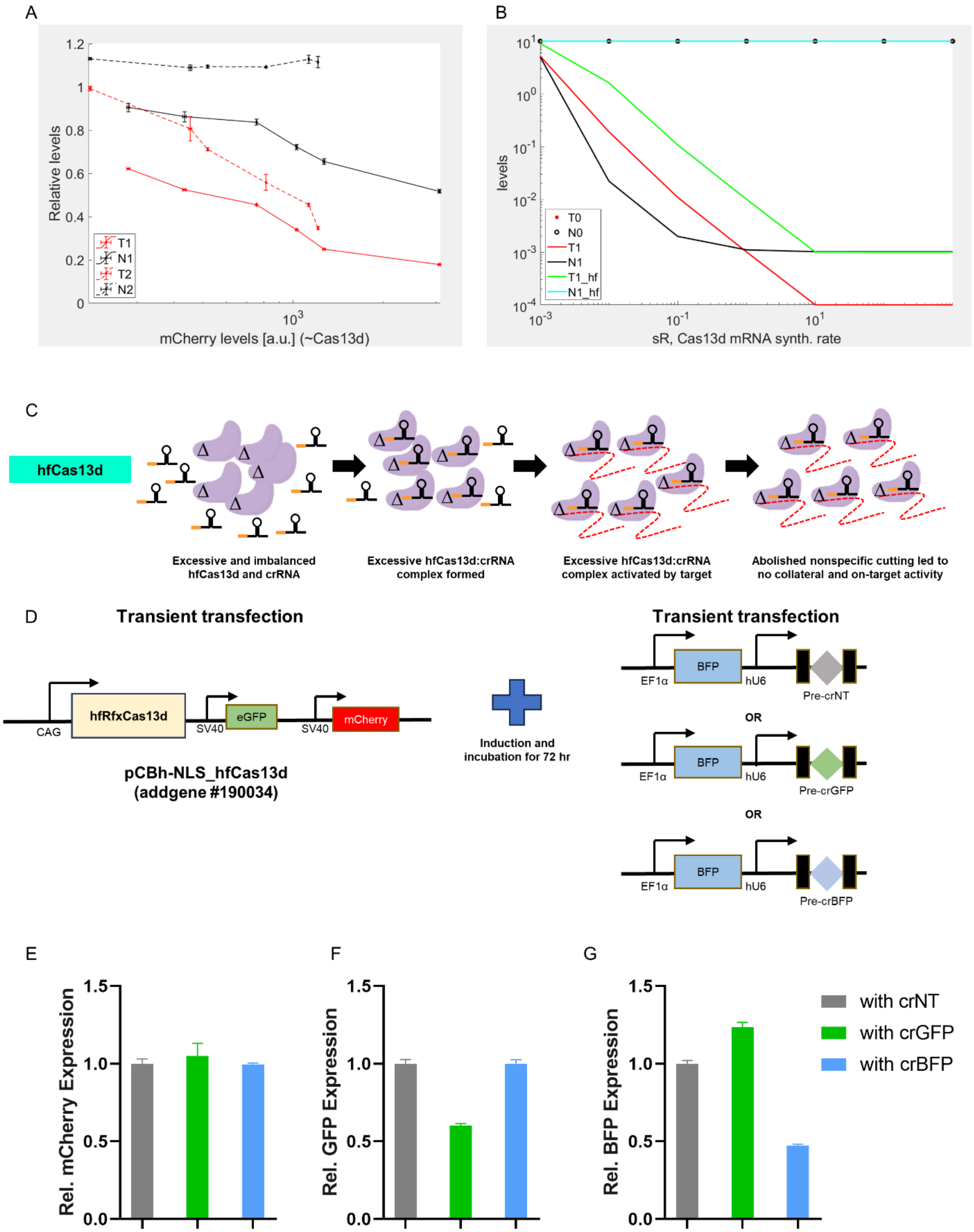
A
Relative levels
mCherry levels [a.u.] (~Cas13d)
T1
N1
T2
N2
B
levels
sR, Cas13d mRNA synth. rate
T0
N0
T1
N1
T1_hf
N1_hf
C
hfCas13d
Excessive and imbalanced hfCas13d and crRNA
Excessive hfCas13d:crRNA complex formed
Excessive hfCas13d:crRNA complex activated by target
Abolished nonspecific cutting led to no collateral and on-target activity
D
Transient transfection
CAG
hfRfxCas13d
SV40
eGFP
SV40
mCherry
pCBh-NLS_hfCas13d (addgene #190034)
Induction and incubation for 72 hr
Transient transfection
EF1α
BFP
hU6
Pre-crNT
OR
Pre-crGFP
OR
Pre-crBFP
E
Rel. mCherry Expression
F
Rel. GFP Expression
G
Rel. BFP Expression
with crNT
with crGFP
with crBFP

**Figure S6. Mathematical modeling and experimental validation indicated on-target efficacy benefits from nonspecific RNA cutting of Cas13d.**

**(A)** Experimental plot of eGFP levels (red lines, corresponding to target RNAs) and BFP levels (black lines, corresponding to nonspecific RNAs) at different mCherry levels (proportional to Cas13d levels) for both the mNF-Cas13d system (solid lines) and MONARCH_2.0 system (dashed lines) (n=3). eGFP and BFP levels were normalized based on crNT constructs for the mNF system, and basal levels for the MONARCH_2.0 system.

**(B)** Simulated plot of mNF-hfCas13d (with original mNF-Cas13d for comparison). *T0* (red circles): no Cas13d basal control (i.e. only pure synthesis and degradation). *N0* (black circles): no Cas13d basal control (i.e. only pure synthesis and degradation). *T1* (red line): mNF-Cas13d system target RNA cutting. *N1* (black line): mNF-Cas13d system nonspecific RNA cutting. *T1_hf* (green line): mNF-hfCas13d system target RNA cutting. *N1_hf* (cyan line): mNF-hfCas13d system nonspecific RNA cutting.

**(C)** Illustration of hypothesized mechanism of experimental observations from high-fidelity RfxCas13d (hfCas13d). Briefly, with hfCas13d, *trans* cleavage is near completely abolished, therefore it will have no collateral activity but at the same time no target RNA degradation from nonspecific RNA cutting, resulting in a lowered on-target efficacy. Red dash curve, degraded target RNA; Grey dash curve, degraded non-target RNA; "Δ" indicates abolished *trans* cleavage.

**(D)** Diagram to test collateral activity and on-target efficacy of hfRfxCas13d with transient fluorescence reporters. Individual crRNAs targeting GFP, BFP or scramble are transfected with hfCas13d expression plasmid together. Flow cytometry examination of each reporter was performed after 72 hours incubation post-transfection.

**(E) (F) (G)** Mean fluorescence intensity of mCherry **(E)**, GFP **(F)** and BFP **(G)** reporter changes after co-transfection of hfCas13d and corresponding crRNA in HEK 293 cells (n=3).

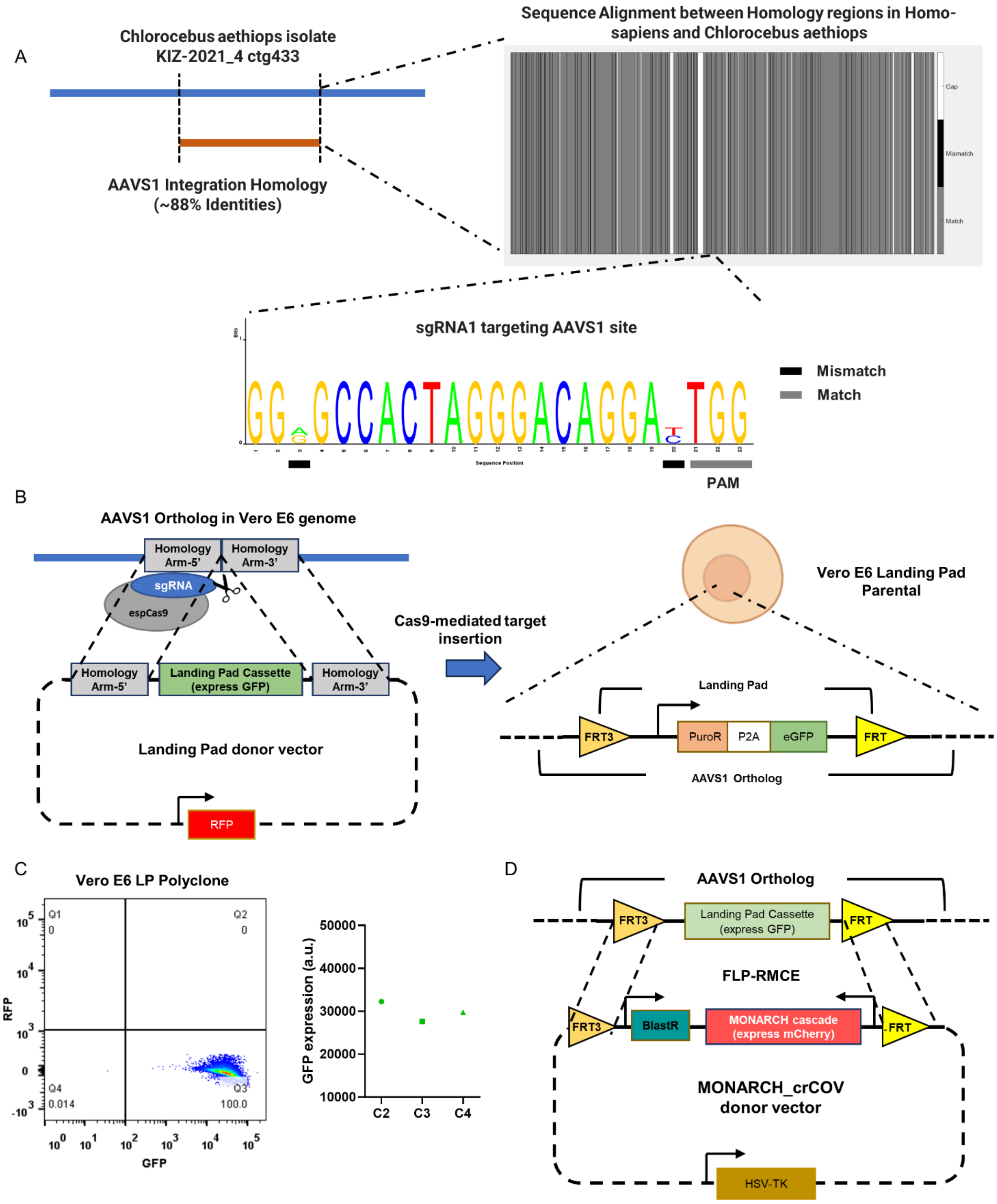
A
Chlorocebus aethiops isolate KIZ-2021_4 ctg433
AAVS1 Integration Homology (~88% Identities)
Sequence Alignment between Homology regions in Homo-sapiens and Chlorocebus aethiops
Gap
Mismatch
Match
sgRNA1 targeting AAVS1 site
Mismatch
Match
Sequence Position
PAM
B
AAVS1 Ortholog in Vero E6 genome
Homology Arm-5'
Homology Arm-3'
sgRNA
espCas9
Landing Pad Cassette (express GFP)
Landing Pad donor vector
RFP
Cas9-mediated target insertion
Vero E6 Landing Pad Parental
Landing Pad
FRT3
PuroR
P2A
eGFP
FRT
AAVS1 Ortholog
C
Vero E6 LP Polyclone
Q1 0
Q2 0
Q4 0.014
Q3 100.0
RFP
GFP
GFP expression (a.u.)
50000
40000
30000
20000
10000
C2
C3
C4
D
AAVS1 Ortholog
FRT3
Landing Pad Cassette (express GFP)
FRT
FLP-RMCE
BlastR
MONARCH cascade (express mCherry)
MONARCH_crCOV donor vector
HSV-TK

**Figure S7. AAVS1 site-specific integration strategy for human cell is compatible with Vero cell engineering.**

**(A)** Sequence BLAST identifies AAVS1 integration homology ortholog in Chlorocebus aethiops with 88% matching identities. Specifically, the PAM site for AAVS1 targeting single-crRNA is valid and only two mismatches are found in the crRNA region.

**(B)** Integrating Landing Pad cassette into Vero E6 cell with the same AAVS1 site-specific insertion strategy using eSpCas9:sgRNA complex developed for human cell. Successful site-specific integration will generate green fluorescence only cells with puromycin resistance.

**(C)** Fluorescence examination of bulk and single-cell sorted Vero E6 clones with successful Landing Pad integration. GFP expression levels were stable and conserved between clones picked and no RFP expression was detected.

**(D)** The similarly applicable RMCE-based DNA constructs switching was used to integrate MONARCH_crCOV cascade into AAVS1 ortholog in Vero E6 landing pad parental cells.

### Supplemental Tables

| Cascade Name | Cas13d promoter | CRISPR RNA | CRISPR RNA promoter | On-target RNA | On-target RNA promoter | Off-target RNA | Off-target RNA promoter |
|---|---|---|---|---|---|---|---|
| mNF-Cas13d | CMV-d2i | N/A | N/A | N/A | N/A | N/A | N/A |
| CMV-Cas13d | CMV | N/A | N/A | N/A | N/A | N/A | N/A |
| mNF-Cas13d_crGFP_eGFP testing platform | CMV-d2i | crGFP | hU6-2O | GFP | SV40 | N/A | N/A |
| mNF-Cas13d_crNT_eGFP control platform | CMV-d2i | crNT | hU6 | EGFP | SV40 | N/A | N/A |
| mNF-Cas13d_crGFP_eGFP_BFP testing platform | CMV-d2i | crGFP | hU6-2O | EGFP | SV40 | BFP | SV40 |
| mNF-Cas13d_crNT_eGFP_BFP control platform | CMV-d2i | crNT | hU6 | EGFP | SV40 | BFP | SV40 |
| eGFP basal control | N/A | N/A | N/A | EGFP | SV40 | N/A | N/A |
| eGFP_BFP basal control | N/A | N/A | N/A | EGFP | SV40 | BFP | SV40 |
| MONARCH 1.0_crGFP | CMV-d2i | crGFP | CMV-d2i with Cas13d | N/A | N/A | N/A | N/A |
| MONARCH 1.0_crGFP_eGFP testing platform | CMV-d2i | crGFP | CMV-d2i with Cas13d | EGFP | SV40 | N/A | N/A |
| MONARCH 1.0_crGFP_eGFP_BFP testing platform | CMV-d2i | crGFP | CMV-d2i with Cas13d | EGFP | SV40 | BFP | SV40 |
| MONARCH 2.0_crGFP | CMV-d2i | crGFP | CMV-d2i with Cas13d | N/A | N/A | N/A | N/A |
| MONARCH 2.0_crGFP_eGFP testing platform | CMV-d2i | crGFP | CMV-d2i with Cas13d | EGFP | SV40 | N/A | N/A |
| MONARCH 2.0_crGFP_eGFP_BFP testing platform | CMV-d2i | crGFP | CMV-d2i with Cas13d | EGFP | SV40 | BFP | SV40 |
| MONARCH 1.0_crCOV | CMV-d2i | crCOV20,21,24 | CMV-d2i with Cas13d | N/A | N/A | N/A | N/A |
| MONARCH 2.0_crCOV | CMV-d2i | crCOV20,21,24 | CMV-d2i with Cas13d | N/A | N/A | N/A | N/A |

**Table S1: List of all the key genetic components of individual circuit and testing cascade stably integrated in the AAVS1 site.**

| Single-guide RNA | Sequence (5’ -> 3’) | Source |
|---|---|---|
| AAVS1-gRNA | GGGGCCACTAGGGACAGGAT | (Wan, Cohen et al. 2023) |
| **CRISPR RNA** | **Sequence (5’ -> 3’)** | **Source** |
| Non-targeting crRNA | GTAACTACTCGAGACGCCTATT | (Abbott, Dhamdhere et al. 2020) |
| BFP-targeting crRNA | TTGAAGTAAAGTTTACGCCTCGA | This study |
| GFP-targeting crRNA | CATGATATAGACGTTGTGGCTGT | This study |
| BACH1-targeting crRNA1 | GTGGTGGGATCTCAGATTTCTGT | This study |
| BACH1-targeting crRNA2 | CTGTCTCGAAATGATTTTCAGTC | This study |
| BACH1-targeting crRNA3 | CTCTGGCCTTGCCTTCTTTATGC | This study |
| SARs-CoV-2-targeting crRNA20 | CACTATTAGCATAAGCAGTTGT | (Abbott, Dhamdhere et al. 2020) |
| SARs-CoV-2-targeting crRNA21 | TTGAATCTGAGGGTCCACCAAA | (Abbott, Dhamdhere et al. 2020) |
| SARs-CoV-2-targeting crRNA24 | AACGCCTTGTCCTCGAGGGAAT | (Abbott, Dhamdhere et al. 2020) |

**Table S2: List of guide RNAs used for AAVS1 CRISPR-Cas9 targeting and Cas13d RNA targeting.** All the crRNAs were designed using web-based online tools, cas13design (https://cas13design.nygenome.org/), developed by Sanjana Lab at NYU (Wessels, Mendez-Mancilla et al. 2020). crRNA candidates with the highest scores were picked for testing.
